# Spatial omics of acute myocardial infarction reveals a novel mode of immune cell infiltration

**DOI:** 10.1101/2024.05.20.594955

**Authors:** Florian Wünnemann, Florian Sicklinger, Kresimir Bestak, Jose Nimo, Tobias Thiemann, Junedh Amrute, Mathias Nordbeck, Niklas Hartmann, Miguel A. Ibarra-Arellano, Jovan Tanevski, Clara Heine, Norbert Frey, Kory J. Lavine, Fabian Coscia, Julio Saez-Rodriguez, Florian Leuschner, Denis Schapiro

**Author notes:** These authors contributed equally (Florian Wünnemann, Florian Sicklinger). These authors jointly supervised this work (Florian Leuschner, Denis Schapiro).

## Abstract

Myocardial infarction (MI) continues to be a leading cause of death worldwide. Even though it is well-established that the complex interplay between different cell types determines the overall healing response after MI, the precise changes in the tissue architecture are still poorly understood. Here we generated an integrative cellular map of the acute phase after murine MI using a combination of imaging-based transcriptomics (Molecular Cartography) and antibody-based highly multiplexed imaging (Sequential Immunofluorescence), which enabled us to evaluate cell-type compositions and changes at subcellular resolution over time. One striking finding of these analyses was the identification of a novel mode of leukocyte accumulation to the infarcted heart via the endocardium - the inner layer of the heart. To investigate the underlying mechanisms driving this previously unknown infiltration route, we performed unbiased spatial proteomic analysis using Deep Visual Proteomics (DVP). When comparing endocardial cells of homeostatic hearts and infarcted hearts, DVP identified von Willebrand Factor (vWF) as an upregulated mediator of inflammation 24 hours post-MI. To further explore the immune mediating capabilities of vWF and its effect on tissue repair, we performed functional blocking of vWF during acute murine MI. This resulted in a reduced amount of infiltration by CCR2^+^ monocytes and worse cardiac function post-MI. Our study provides the first spatial map of acute murine MI with subcellular resolution and subsequently discovers a novel route of immune infiltration. Furthermore, we identified vWF as a critical immune mediating agent for endocardial immune cell infiltration.

## Main

Myocardial infarction (MI), also known as a heart attack, is an acute disease characterized by large shifts in cellular composition and tissue architecture due to cell death of cardiac muscle tissue caused by local hypoxia ^1^. MI remains one of the leading causes of death worldwide, despite large improvements in the prevention and treatment of the disease ^2,3^. While advancements in restoring blood flow to the heart muscle (termed reperfusion) and pharmacological treatment strategies have largely reduced short-term deaths, long-term mortality after MI continues to be high ^4–6^. One of the reasons why mortality remains high is that we do not understand how molecular cues and tissue microenvironment alterations during acute disease affect healing and remodeling in the long run. During an acute infarct, necrotic cells in the heart release stress signals, including pro-inflammatory cytokines and chemokines, leading to the invasion of the infarct zone by immune cells - specifically neutrophils, monocytes/macrophages (Mo/Mϕ) and later T-cells, B-cells and natural killer cells ^1,7–10^. Modulation of the types and amount of immune cells infiltrating the infarct have been postulated as potential treatment targets to improve healing and outcome after MI ^11,12^. A better understanding of the immune cell infiltration routes and pathways, detailed tissue microenvironment and cellular interactions in the heart during the early phases of MI thus holds the promise to deliver potential novel treatment strategies.

Extensive research has been performed on the temporal dynamics of immune cell infiltration during the course of MI in humans and mice ^1,8,13,14^. Cell count estimations of the immune infiltrate have been generated using Fluorescence-activated Cell Sorting (FACS) of left ventricular cardiac tissue ^14–17^. Single-cell RNA sequencing (scRNA-seq) provided further insight into the diverse sub-types of immune cells that infiltrate the necrotic myocardium and their different pathway activities and physiological roles ^18–21^. Two recent studies applied single-cell nucleus sequencing (snRNA-seq) alongside untargeted spatial transcriptomics to investigate the border infarct zone during MI in mice ^22,23^. Combining the spatial readout with transcriptome-wide measurements, both studies identified important transcriptional signatures of the infarct border zone. Another large study generating a spatial multi-omic map of human MI also utilized untargeted transcriptomics to investigate the different cell neighbourhoods and tissue architectures during MI progression in humans ^24^. While these studies were able to connect critical molecular patterns and processes during MI to their spatial context, they could not accurately quantify the tissue microenvironment and cellular neighbourhoods, due to the limited spatial resolution (∼55µm) of the technology used, which effectively captures a mix of 10-20 cells per measurement. Novel targeted spatial omics technologies with subcellular resolution are transforming our understanding of tissue architecture and the corresponding cell-type interactions in health and disease ^25,26^. Highly multiplexed approaches to measure tens to thousands of transcripts, antibodies or both combined, enable a detailed description of the changing cellular phenotypes and neighbourhoods during homeostasis and disease ^27–30^. Furthermore, developments in computer vision now enable cell identification using automated cell segmentation and classification algorithms ^31–34^.

Here, we utilized combinatorial imaging technologies based on RNA detection via fluorescent in-situ hybridization (FISH) barcoding (Molecular Cartography™) and antibody-based, Sequential Immunofluorescence (SeqIF, Lunaphore COMET™), respectively, to characterize the changing tissue microenvironment during acute MI in mice. Subsequently, we developed novel, scalable computational pipelines (nf-core/molkart) tailored for cardiac tissues, utilizing state-of-the-art methodology to process these complex, highly-multiplexed imaging data sets. Using Molecular Cartography and SeqIF across a time course of the acutely infarcted heart (control prior to infarct, 4 hours, 24 hours, 2 days, 4 days post-MI), in a minimally invasive MI model ^35^, we were able to characterize the acute MI microenvironment at single-cell resolution. From this spatiotemporal map of MI, we discovered that myeloid cells (specifically Mo/Mϕ) enter the infarct region not only via the border and epicardial infarct zone, but also via the endocardium, which was previously unknown. Laser microdissection of endocardial regions 24 hours after infarction followed by ultrasensitive proteomics ^36^ was performed to investigate this novel infiltration route, which revealed local signatures of inflammation and coagulation factors and highlighted von Willebrand Factor (vWF) as an immune mediating agent in endocardial cells. To further investigate the role of vWF, antibody-based functional blocking of vWF during the first day following MI showed a significant reduction of Mo/Mϕ infiltration via the subendocardial infarct zone and reduced left ventricular function. Therefore, our study highlights, for the first time, a critical role of the endocardium for infiltration of immune cells into the infarct via local upregulation of adhesion factors such as vWF. These results highlight previously unknown routes of immune cell infiltration and provide novel potential targets for pharmacological intervention.

### Single cell resolved spatial transcriptomic analysis of acute MI

To characterize the cellular environment during homeostasis and acute MI in the mouse heart, we used a combinatorial single-molecule FISH (smFISH) based technology called Molecular Cartography by Resolve Bioscience (**Figure 1a, Supplementary Figure 1a-c**). Molecular Cartography allows for the detection of RNA transcripts for up to 100 candidate genes at single molecule resolution with high sensitivity in selected regions of interest (ROI). Based on marker gene expression from existing publicly available single-cell RNA-seq datasets and expert knowledge ^37–41^, we subsequently designed a 100 gene transcript panel specifically for this study, to capture major cell types as well as inflammatory signals during acute MI (**Supplementary Table 1**). Using this panel, we selected one ROI per heart (2.09 mm^2^ to 2.97 mm^2^) cross-section to measure endocardial, myocardial and epicardial regions at 4 different time points during acute MI (control prior to infarct, 4 hours, 2 days, 4 days) (**Supplementary Figure 1a-c**). Identification and abundance of RNA transcript spots were highly reproducible across technical replicate slides of Molecular Cartography, as highlighted by a strong correlation of bulk transcript counts across sections from the same biological sample (**Supplementary Figure 2a**). On average across all time points, we detected 980 000 transcript spots per mm^2^ of heart tissue with lower transcript density in infarcted tissues due to less viable cells within the infarct region (**Supplementary Figure 2b**). Principal component analysis (PCA) of pseudo-bulk transcriptional profiles showed separation of samples by time relative to the induced infarct, confirming strong transcriptional shifts in response to acute infarction (**Supplementary Figure 2c**). Molecular Cartography data was processed with an in-house developed computational pipeline which we call nf-core/molkart (**Figure 1b, Supplementary Figure 3**). nf-core/molkart will be available open-source with this manuscript. Despite advances in deep learning-based methods, segmentation of cardiac cells from microscopy images remains extremely challenging due to their differences in cell size, shape, orientation, multinucleation and RNA content. Comparing three different cell / tissue segmentation methods (Mesmer Deepcell ^33^, Ilastik Multicut ^42^ (not shown), Cellpose 2 ^34^), we found that Cellpose 2 with a human-in-the-loop trained model performed best on cardiac tissues when evaluating cell shapes, cell sizes and the percent of assigned RNA spots overall (**Supplementary Figure 4a,b**). Cell typing of cell transcript profiles across all images (69028 cells in total, average number of cells per sample = 8629) identified all major cardiac cell types during acute MI (**Figure 1c**). We found healthy as well as stressed cardiomyocytes distinguished by their expression of atrial natriuretic peptide (Nppa) and brain natriuretic peptide (Nppb), cells of the vasculature (pericytes, smooth muscle cells, endothelial cells), cardiac fibroblasts as well as infiltrating myeloid and lymphoid cells (**Figure 1d**). The hypoxic environment in the left ventricle (LV) during acute MI leads to massive cell death and changes in tissue composition and architecture within the LV. In line with these expectations and known cell dynamics from literature, we observed a strong shift in the cell type composition from healthy to infarcted LV tissue during the first 4 days after MI (**Supplementary Figure 5a,b**). Healthy LV tissue composed of cardiomyocytes, endothelial cells and fibroblasts showed compositional changes towards an environment of dying and stressed cardiomyocytes (Nppa^+^) surrounded by extracellular matrix (ECM) producing cardiac fibroblasts and invading myeloid cells after MI. Since our selected regions of interest (ROI) contained large areas of infarct tissue, healthy cardiomyocytes showed the highest decrease in cell number from an average of 57.9% of identified cells in controls to only about 10% at day 4 in the processed regions. In line with increased ECM production and early scar formation, cardiac fibroblasts increased from 6.6% to 30.8 % of cells within the ROIs. Notably, we observed a strong increase of myeloid cells (neutrophils, Mo/Mϕ and dendritic cells) from 4% in controls to almost 28% at day 4 in the imaged ROIs, indicating ubiquitous infiltration of myeloid cells into the infarct (**Supplementary Figure 5b**). Overall, our analysis highlights the cellular landscape of acute MI, recapitulating many known cellular dynamics in a spatial context.

**Figure 1:**
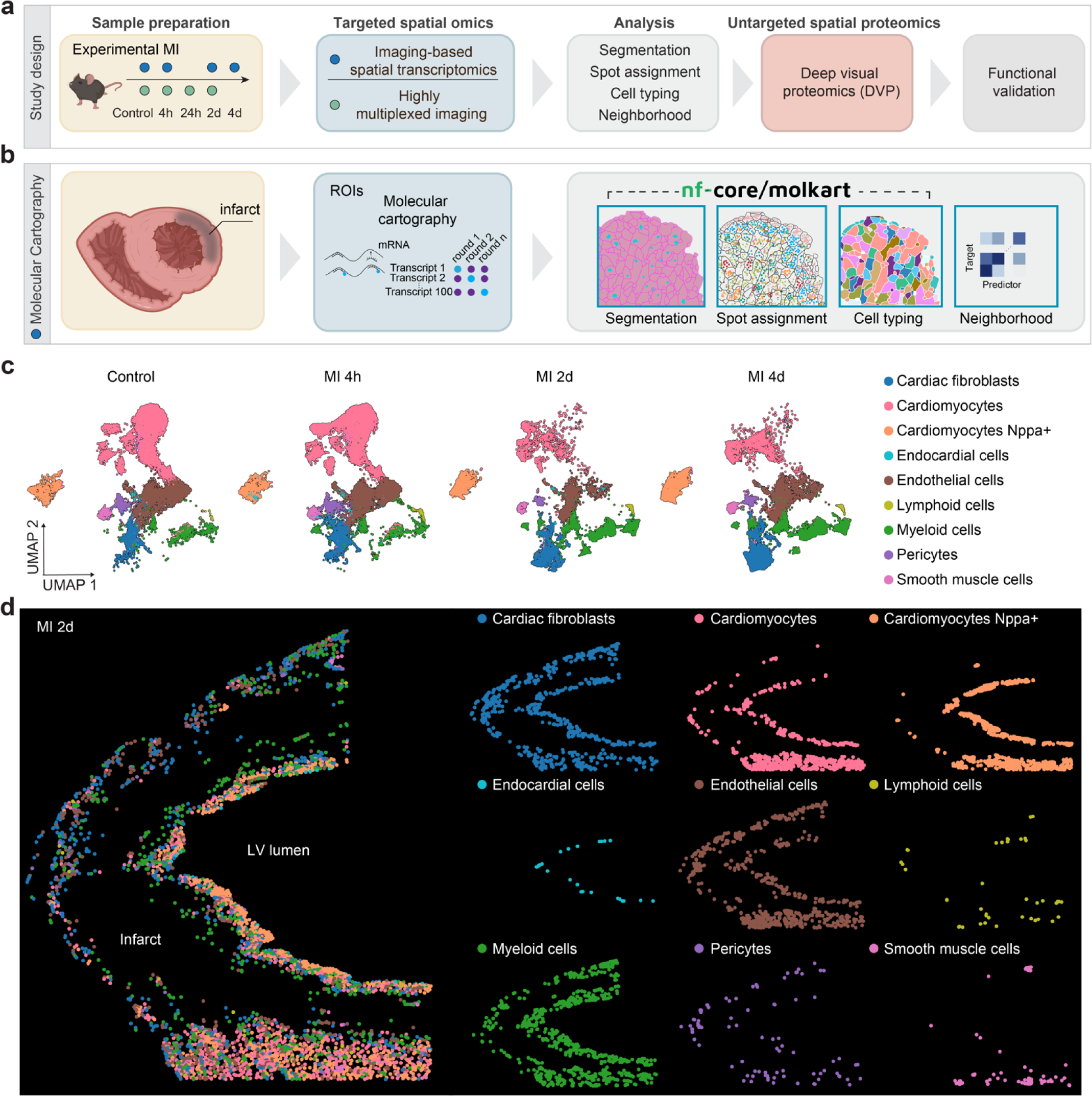
Molecular Cartography of acute myocardial infarction enables spatial cell-typing of the left ventricular infarct tissue. **a)** Schematic overview of the study design. **b)** Schematic for spatial transcriptomics data generation (Molecular Cartography) and processing (nf-core/molkart). **c)** UMAP showing the joint embedding of 69028 cells from 8 samples over four time points across acute MI. **d)** Representative spatial cell-type distributions for a sample at 2 days post-MI. A composite image with all cell types is shown on the left, while each cell type’s individual distribution is shown on the right.

### Cellular neighbourhood analysis of spatial transcriptomic data highlights the dynamic spatiotemporal changes during acute MI

To get a global understanding of the tissue architecture and intercellular relationships across acute MI tissues, we applied MISTy (Multiview Intercellular SpaTial modeling framework) ^43^ to our dataset. MISTy captures the cell type relationship patterns across entire slides and datasets in an unbiased manner (Methods). In homeostatic cardiac control tissue, the majority of cell types were distributed mostly homogeneously, with cardiomyocytes (CM) interspersed with cardiac fibroblasts (CF), vascular endothelial cells and pericytes. The location of the majority of cell types therefore was not informative as a predictor of the localization of other cell types. MISTy did however identify a spatial signature for Nppa^+^ cardiomyocytes, whose spatial localization was best predicted by endocardial cells (**Figure 2a**). Myeloid cells in control tissue were rare with a relatively homogeneous distribution across the tissue during homeostasis and 4 hours after MI, indicating that these are likely resident leukocytes. To further explore the potential colocalization and interaction between cell types, we performed local, bi-variate analysis between pairs of cell types using Liana+ ^44^. This local analysis between endocardial cells and Nppa^+^ cardiomyocytes in controls pinpointed the interaction between these cell types to the subendocardial region close to the LV lumen (**Figure 2b**). This aligns well with known localization of Nppa in trabecular ventricle regions during development, which is maintained in homeostatic adult hearts ^45–47^. In contrast, no spatial interaction was identified between myeloid cells and endocardial cells in controls (**Figure 2c**). Visualization of the RNA signal within that region confirmed strong Nppa expression close to markers for endocardial / endothelial cells (Pecam1) (**Figure 2d**). Two days after MI, MISTy analysis revealed interactions that were not identified in control conditions. Besides the remaining strong relationship between endocardial cells and Nppa^+^ cardiomyocytes, MISTy also identified a new spatial context between endocardial cells and myeloid cells (**Figure 2e**). Both of these spatial interactions showed clear demarcation of a (sub)endocardial infarct zone around the infarct core (**Figure 2f,g**). In line with this finding, we observed a strong RNA signal of myeloid markers within the endocardium and the subendocardial infarct zone (demarcated by Nppa) (**Figure 2h**). The local relationship between endocardial cells and myeloid cells was also identified at 4 days post-MI alongside a new signal between CF and myeloid cells (**Figure 2i**). Myeloid cell locations at 4 days post-MI were additionally predicted by Nppa+ cardiomyocytes, cardiac fibroblasts and cardiomyocytes (**Figure 2j**). Interestingly, the interaction between myeloid cells and fibroblasts was enriched in the border zone and epicardial infarct zone (**Figure 2k**). The highly increased abundance of CF and their spatial colocalization with myeloid cells were further highlighted by the drastic increase of RNA molecules encoding extracellular matrix components like Col1a1 (**Figure 2l**). As multiple spatial analyses highlighted unexpected but potentially important interactions between endocardial cells and myeloid cells, we aimed to validate and quantify this increased local relationship using simple measures. Therefore, we calculated the Euclidean distance in two-dimensional tissue space between each endocardial cell and its three nearest neighbour myeloid cells to quantify myeloid cell proximity to endocardial cells during acute MI (**Figure 2m**). Average distances between endocardial cells and myeloid cells showed significant differences over the MI time course (**Figure 2n**). At day 2 and day 4 after MI, endocardial cells showed significantly shorter distances to myeloid cells, while the average distance to myeloid cells did not change significantly between day 2 and day 4 (**Figure 2n**). We performed the same distance analysis to see whether myeloid cells show a similar relationship with CF at the interface to the infarct core and found increased proximity between both cell types after MI (**Figure 2o**). Taken together, our cellular neighbourhood analysis identified and highlighted an unexpected spatial relationship between endocardial cells and myeloid cells, suggesting that immune infiltration to the infarct might be mediated via the endocardium and the subendocardial, Nppa^+^ positive infarct zone.

**Figure 2:**
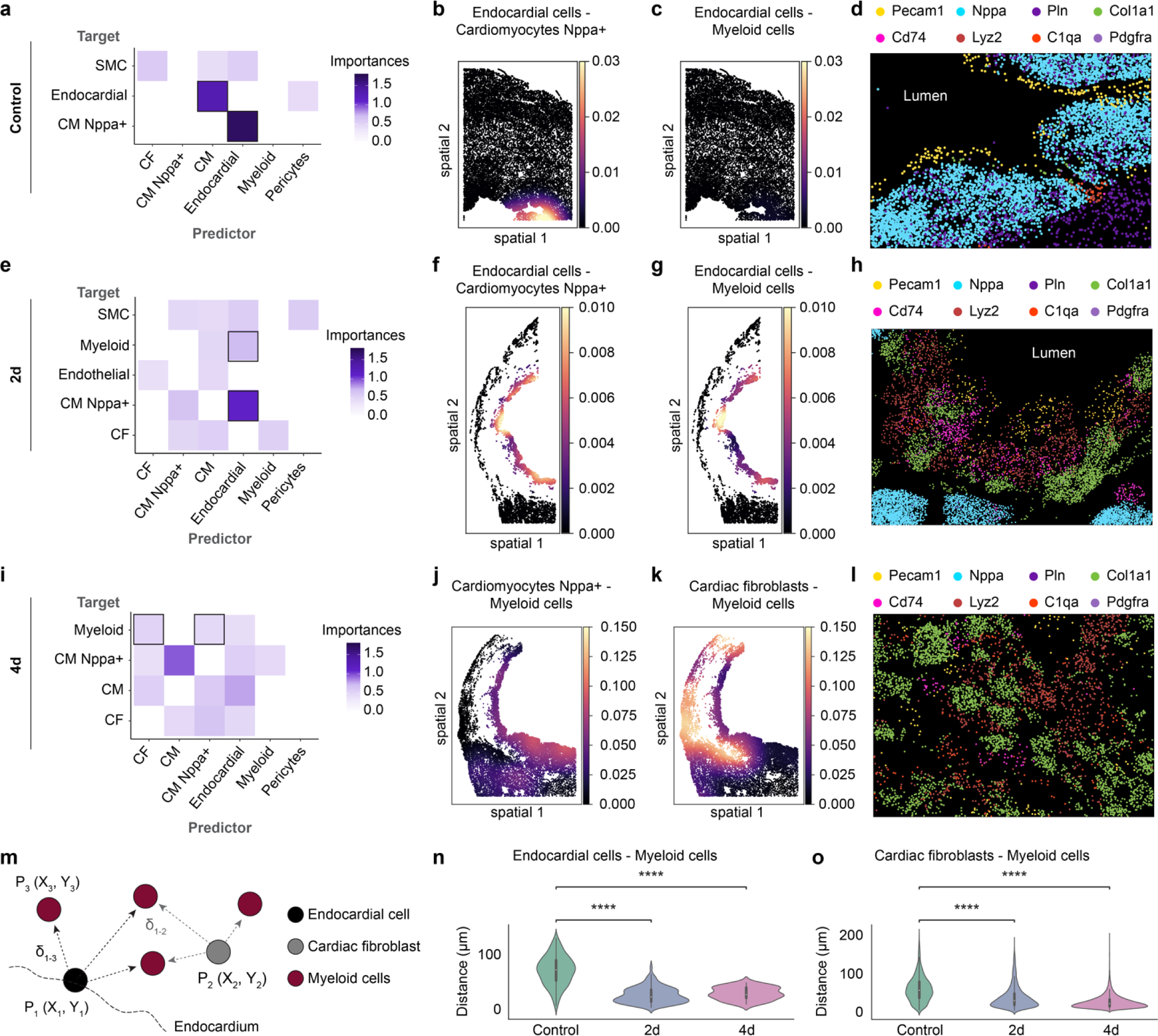
Spatial analysis of Molecular Cartography cell composition during MI highlights myeloid interactions with the endocardial layer. Abbreviations in the figures represent CM = cardiomyocytes, CF = cardiac fibroblasts and SMC = smooth muscle cells. **a)** Spatial cell type relationships in cardiac control tissue as calculated by MISTy. Importance indicates spatial interactions across the slide between the two cell types highlighted. **b), c)** Local-bivariate analysis between endocardial cells and cardiomyocytes Nppa^+^ and endocardial cells and myeloid cells, respectively. Colour indicates the local product as calculated by Liana+. **d)** RNA spot localization in an endocardial region of control tissue, highlighting spatial co-localization of marker genes for endothelial/endocardial cells (Pecam1), Cardiomyocytes (Pln, Nppa), Fibroblasts (Pdgfra, Col1a1) and myeloid cells (Cd74, Lyz2, C1qa). **e)** MISTy analysis for LV tissue 2 days post-MI shows a new interaction between endocardial cells and myeloid cells. **f), g)** Local interaction analysis shows the interaction between endocardial cells and myeloid cells in the subendocardial infarct zone, also marked by Nppa. **h)** RNA marker expression confirming localized expression of myeloid markers in the subendocardial infarct zone. **i)** MISTy analysis for 4 days post-MI highlights the spatial relationship between cardiac fibroblast and myeloid cells around the infarct core. **j), k)** Local analysis highlighting cardiac fibroblast, cardiomyocytes Nppa+ and myeloid interactions. **l)** RNA spot localization within the infarct tissue at 4 days post-MI. **m)** Euclidean distances between all pairs of cell types were calculated. The three closest neighbours per cell were taken as an estimate of average distances between cell types. **n)** Euclidean distances between endocardial cells to myocardial cells were significantly different across the four first days after MI (Type II ANOVA p-value = 6.54e-88). Post-hoc analysis showed significant differences at 2 days (post-hoc t-test with Bonferroni correction, p-value=4.04e-78) and 4 days (p-value=8.28e-22) post-MI relative to control but no difference between 2 days and 4 days (p-value = 0.18). **o)** Euclidean distances between cardiac fibroblasts and myeloid cells were also significantly different across time. For violin plots, white dots indicate the median, black boxes indicate the interquartile range (IQR) and whiskers represent 1.5 x IQR. For MISTy analysis, only interactions with an importance > 0.4 and only cell types with a gain in R^2^ > 5 are shown.

### Highly multiplexed antibody-based imaging confirms immune cell infiltration via the endocardial layer in acute MI

To further investigate the regional distribution of myeloid cells after MI and capture spatial temporal patterns across entire heart sections, we performed Sequential Immunofluorescence (SeqIF™ ^48^) on samples from the acute phase after MI (control, 4 hours, 24 hours, 2 days) (**Figure 3a**). We optimized an antibody panel to identify healthy and stressed cardiomyocytes (Tnnt2, Ankrd1), endothelial cells (CD31), smooth muscle cells (aSMA), cardiac fibroblasts (Pdgfra), myeloid cells (CD45, CD68, CCR2, Trem2, Mpo), as well as DAPI and WGA to capture nuclei and the cell membrane respectively (**Figure 3b**, **Supplementary Table 2**). We performed image processing of SeqIF data using MCMICRO, a Nextflow-based pipeline that performs subtraction of autofluorescence signal from each antibody channel, cell segmentation and fluorescence intensity quantification ^49^ (**Figure 3a**). Based on our experience with segmentation for Molecular Cartography data, we applied the same strategy of using WGA and DAPI to train a custom Cellpose 2 model to segment cells in the SeqIF dataset. To assign phenotypes to cells, we used the Pixie workflow ^50^, which performs pixel clustering using self-organizing maps (SOMs) to generate pixel maps of tissues (see Methods for more details) (**Supplementary Figure 6a**). In these pixel maps, groups of pixels with similar marker intensity profiles across the SeqIF dataset are clustered together, allowing for classification of different cell types and tissue regions across the entire time course of acute MI (**Supplementary Figure 6b**) ^50^. In line with our results from Molecular Cartography, we found a continuous decrease in pixel phenotype clusters for cardiomyocyte marker Tnnt2, and an increase in pixel phenotype clusters for myeloid cells (Mo/Mϕ and neutrophils) in the first two days after MI (**Supplementary Figure 6b,c**). We also found stressed and dying cardiomyocytes positive for an Ankrd1 pixel cluster, clearly demarcating the infarct core starting already at 4 hours after MI (**Supplementary Figure 6a**). We used the pixel maps to perform cell phenotyping using Cellpose cell masks in a second clustering step (**Figure 3c,d**). This pixel-level phenotyping workflow enabled us to profile the spatial localization of cardiac cells within the infarcted heart based on our highly-multiplexed imaging data, at an unprecedented scale for entire heart cross-sections. To further investigate and independently validate the potential relationship between endocardial and myeloid cells that we found in Molecular Cartography data, we repeated the distance analysis between these two groups of cells. In line with our findings with Molecular Cartography, the distance between myeloid cells and endocardial cells decreases during the first 2 days after MI (**Figure 3e**). Interestingly, myeloid cells were closest to endocardial cells only 24 hours following an infarct (median distance = 24µm), suggesting that attachment and infiltration of these immune cells via the endocardial layer might be a rapid process. To quantify the extent of different infiltration routes of myeloid cells into the infarct, we partitioned the infarcted heart images into regional compartments (endocardium, epicardium, infarct core and border zone) guided by expression patterns of Tnnt2, Ankrd1, WGA and CD31 (see Methods section) (**Figure 3f**, **Supplementary Figure 7a**). Using both, SeqIF and conventional immunofluorescence stainings, we found a strong increase specifically for monocytes/macrophages (Mo/Mϕ) expressing CCR2 (CCR2^+^ Mo/Mϕ) in the endocardium, peaking at 24 hours after MI (**Figure 3g, Supplementary Figure 7b,c**). At 2 days after MI, the relative number of myeloid cells remained high within the endocardial layer. However, we additionally found an increased density of CCR2^+^ Mo/Mϕ within the epicardial infarct layer. Quantification of absolute CCR2^+^ Mo/Mϕ numbers over time identified the border zone as the predominant invasion route after 2 days. Closer inspection of CCR2^+^ Mo/Mϕ distribution in the endocardial layer showed CCR2^+^ Mo/Mϕ, already being attached to the endocardium at 4 hours post-MI, with occasional infiltration events (**Figure 4a-c**). In the epicardial layer, CCR2^+^ Mo/Mϕ were either directly attached to the epicardium or in close proximity to epicardial vessels (**Supplementary Figure 8a-c**). Taken together, our results - across multiple technologies and quantification methods - clearly indicate a progressive infiltration of myeloid cells via different cellular layers during the acute phase after MI in a non-reperfusion context and highlight a novel route via the endocardium that immune cells can take to infiltrate the LV to reach the infarct region.

**Figure 3:**
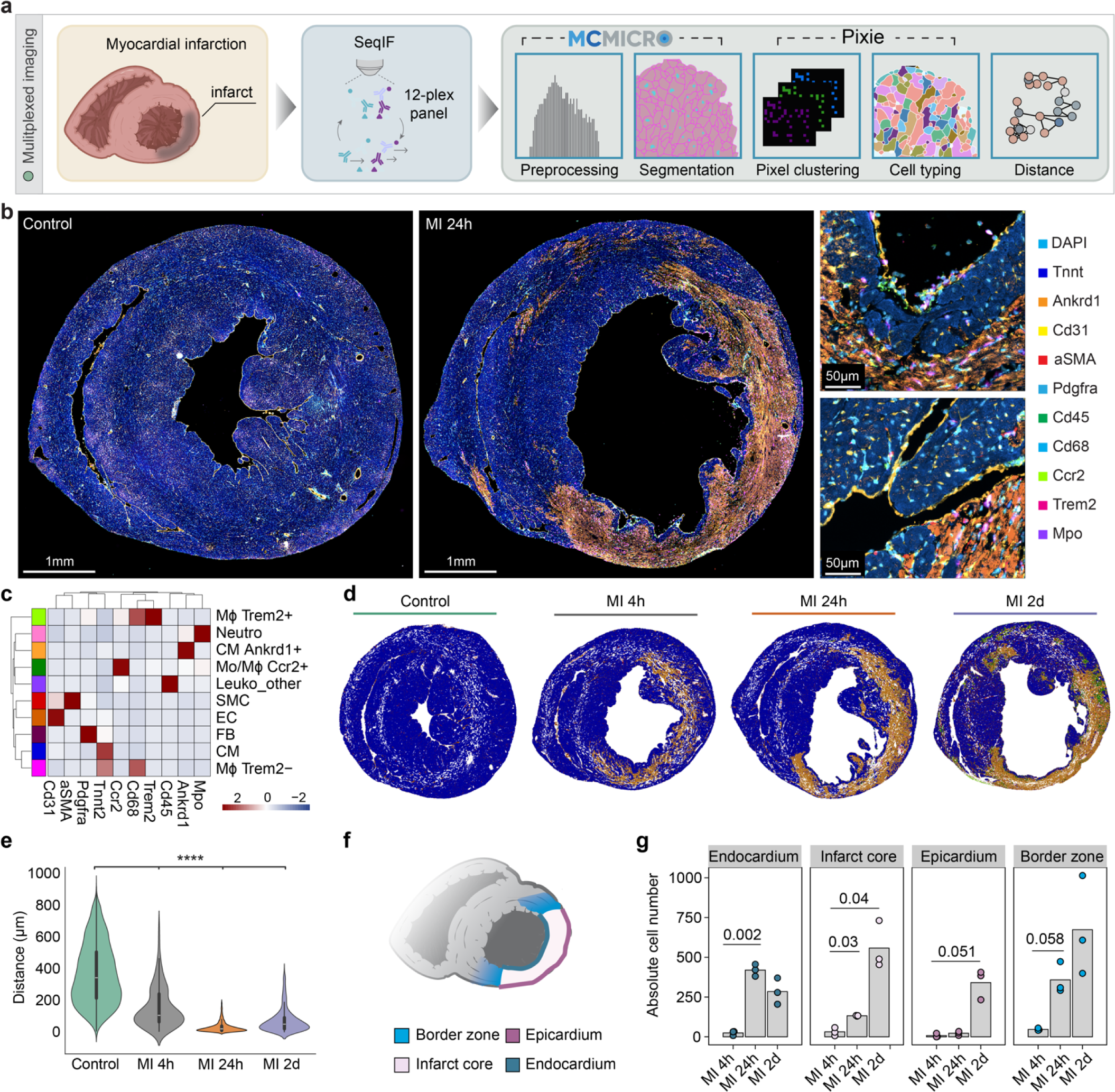
Highly multiplexed imaging using SeqIF and conventional IF during the first two days of acute MI confirms infiltration of myeloid cells via the endocardial layer. **a)** Schema of experimental design for SeqIF data generation and processing. **b)** Sequential immunofluorescence (SeqIF) of mouse heart cross sections using 10 antibodies before (left) and at 24 hours after MI (right). Magnifications on the right highlight endocardial niches in the infarct, characterized by the presence of stressed cardiomyocytes (Ankrd1+, orange) and attachment of immune cells to endocardial cells (CCR2+ in green, Mpo+ in violet and CD31 in yellow). Scale bar full image = 1mm and ROIs = 50µm. **c)** Cell phenotyping heatmap from Pixie highlighting marker expression in different cell masks across the entire dataset. Legend represents the scaled marker expression, which was capped at 3. **d)** Pixie cell phenotyping for one representative sample per time point. Each pixel is colored based on its cell pixel cluster as calculated by Pixie. Cell type colors correspond to heatmap grouping colors in c). **e)** Violin plot of distances in micrometers from endocardial cells to the closest three myeloid cell neighbours quantified at four different time points from SeqIF images. Comparisons between all displayed timepoints show significant differences (post-hoc t-tests with Bonferroni correction for all comparisons p < 1e-16). **f)** Anatomical annotation used to calculate myeloid cell infiltration. For violin plots, white dots indicate the median, black boxes indicate interquartile range (IQR) and whiskers represent 1.5 x IQR. **g)** Quantifications of CCR2+ CD68+ Mo/Mϕ in different spatial regions in cross sections stained with conventional IF. Cell-type abundance is visualized as absolute counts. Bars show mean abundance and points represent individual measurements. *P* values were determined by 2-way ANOVA followed by Tukey’s multiple-comparison test. Only comparisons between timepoints within each region are displayed. CM = cardiomyocytes, Leuko = leukocytes, SMC = smooth muscle cells, EC = endothelial cells, FB = fibroblasts, Mo = monocytes, Mϕ = macrophages.

**Figure 4:**
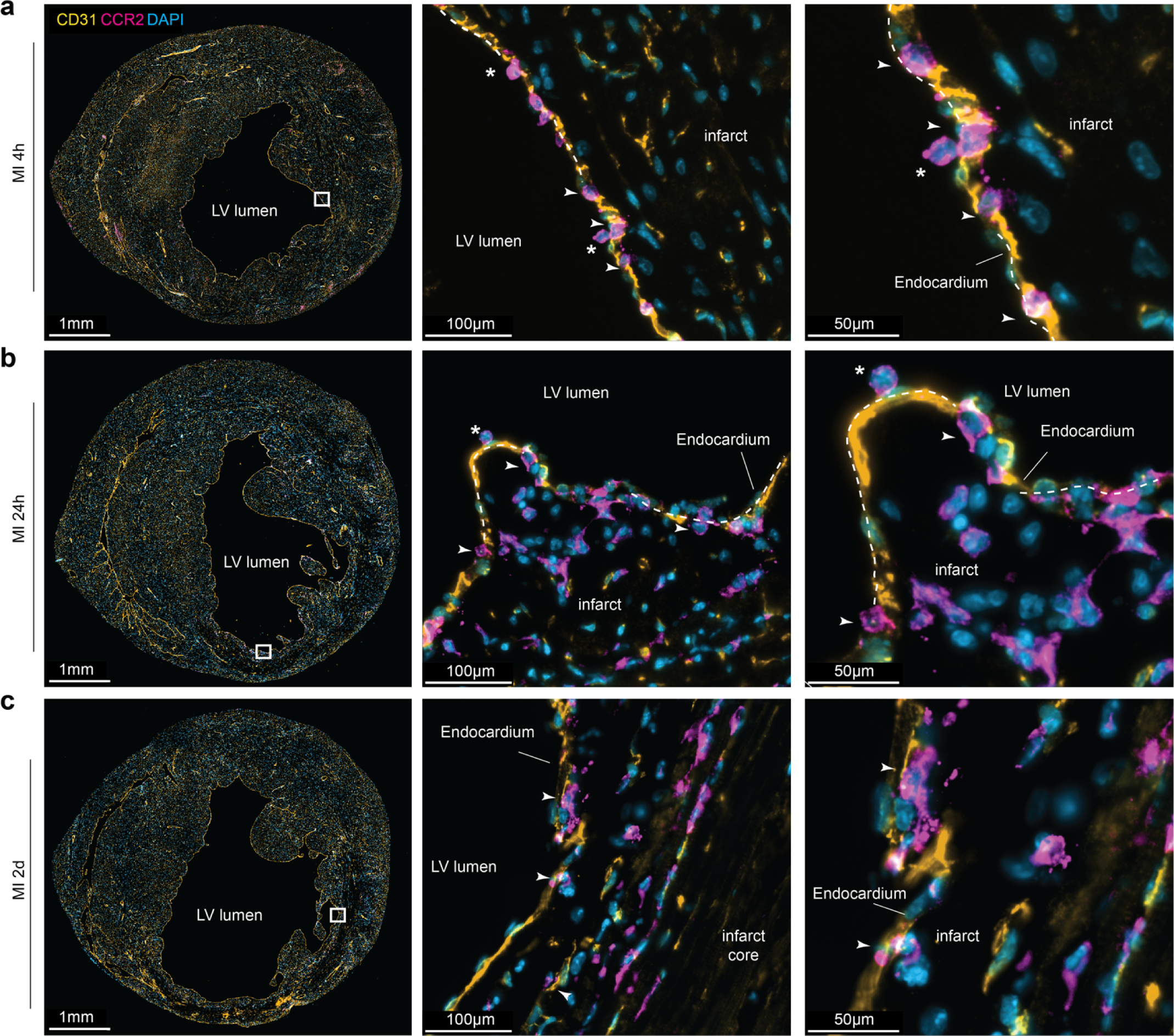
Mo/Mϕ infiltrate the infarcted heart via the endocardium. **a-c)** SeqIF stainings showing selected markers including CD31 (yellow), CCR2 (magenta) and DAPI (blue). Staining of sections 4h (a), 24h (b) and 2 days (c) post-MI indicates increased attachment (asterisk) and transmigration (arrow) resulting in high density of CCR2+ cells in the subendocardial infarct zone and movement of CCR2+ cells towards the infarct core. Mid and right panels represent magnifications of the marked box in the left overview panel.

### Spatial proteomics of the endocardial layer highlights vWF involvement in immune cell infiltration

Following the discovery of early immune cell infiltration into the subendocardial infarct zone, we aimed to identify potential factors mediating the recruitment, adhesion and infiltration of myeloid cells via the endocardium using ultrasensitive mass spectrometry-based proteomics ^36^. Therefore, we utilized laser capture microdissection to excise the endocardial region from healthy mouse hearts (Control) and hearts 24 hours after MI. For hearts with MI, we split endocardial cells into two groups, those that were within the infarct zone (MI IZ) and those that were remote to the infarct (MI remote) (**Figure 5a**). Proteomes of the different endocardial tissue samples encompassed an average number of 3274 proteins after quality control filtering (**Supplementary Figure 9a**). Despite the very low amount of input tissue material, the proteomic data was of high quality with a low amount of missing protein values (4 to 16% across samples) (**Supplementary Figure 9b**). PCA of proteomic samples showed clear separation of control endocardial samples compared to endocardial cells from infarcted samples, indicating a reproducible perturbation of the endocardial layer protein signature 24 hours after MI (**Figure 5b, Supplementary Figure 9c**). Interestingly, the remote endocardial layer signature from infarcted hearts (MI remote) was sufficiently different from control endocardial cells to distinguish them in the PCA, but there were very few significantly differentially expressed proteins (DEPs) between these two conditions (**Supplementary Figure 9d**). In contrast, significant differential protein expression of many proteins was observed in the endocardial region in the infarct (MI IZ) relative to the remote endocardial region (MI remote, **Figure 5c**). Pathway analysis using hallmark gene-sets of DEPs between MI IZ and MI remote revealed upregulated pathways related to immune cell activation (complement system, inflammatory response, interferon-gamma response) as well as downregulated pathways related to energy metabolism (oxidative phosphorylation, fatty acid metabolism) (**Figure 5d**). Interestingly, we identified genes for coagulation pathways strongly upregulated in MI IZ samples relative to MI remote regions (**Figure 5d**). We investigated the cell specificity of these pathway results using a published single-nucleus RNA-seq (snRNA-seq) dataset ^22^ and found vWF as the most specific endocardial protein that was significantly upregulated in MI IZ compared to MI remote, similar to known endothelial adhesion molecules such as Vcam1 (**Figure 5e,f**). vWF is a multimeric protein that plays a central role in vascular homeostasis and is involved in inflammatory processes ^51^. Interestingly, vWF was not significantly upregulated in the remote endocardial regions of infarcted hearts (MI remote) compared to endocardial regions of control hearts (**Figure 5f**). To confirm increased localization of vWF proteins in the endocardial infarct zone, we performed conventional immunofluorescence staining of vWF in infarcted hearts and found a significantly stronger signal in the endocardial infarct area, with an almost absent signal in the remote region (**Figure 5g**). The distribution of vWF positive staining interestingly was not uniform across the infarct adjacent endocardial layer, but stronger at endocardial sites where the ventricular tissues formed pockets, compared to smooth regions. Immunofluorescence stainings of murine hearts 24h after ischemia/reperfusion injury also showed similar staining patterns with increased vWF expression (**Supplementary Figure 10a,b**). To investigate whether vWF also plays a role in human MI, we reprocessed a Cellular Indexing of Transcriptomes and Epitomes by sequencing (CITE-seq) dataset of explanted human hearts from donors and acute MI patients (**Figure 5h**) ^52^. This dataset consists of healthy donors, acute MI (AMI), chronic ischemic and non-ischemic cardiomyopathy patients, from which we focused on healthy donors (n=6) and AMI patients (n=4). Differential gene expression analysis demonstrated a significant increase of endocardial vWF expression in patients after acute MI compared to healthy donors (**Figure 5i**). Collectively, our Deep Visual Proteomics (DVP) analysis during acute MI revealed local spatial differences between endocardial regions within the same heart and highlighted upregulation of vWF as a specific response of the endocardium to the local inflammatory signals from the infarct zone.

**Figure 5:**
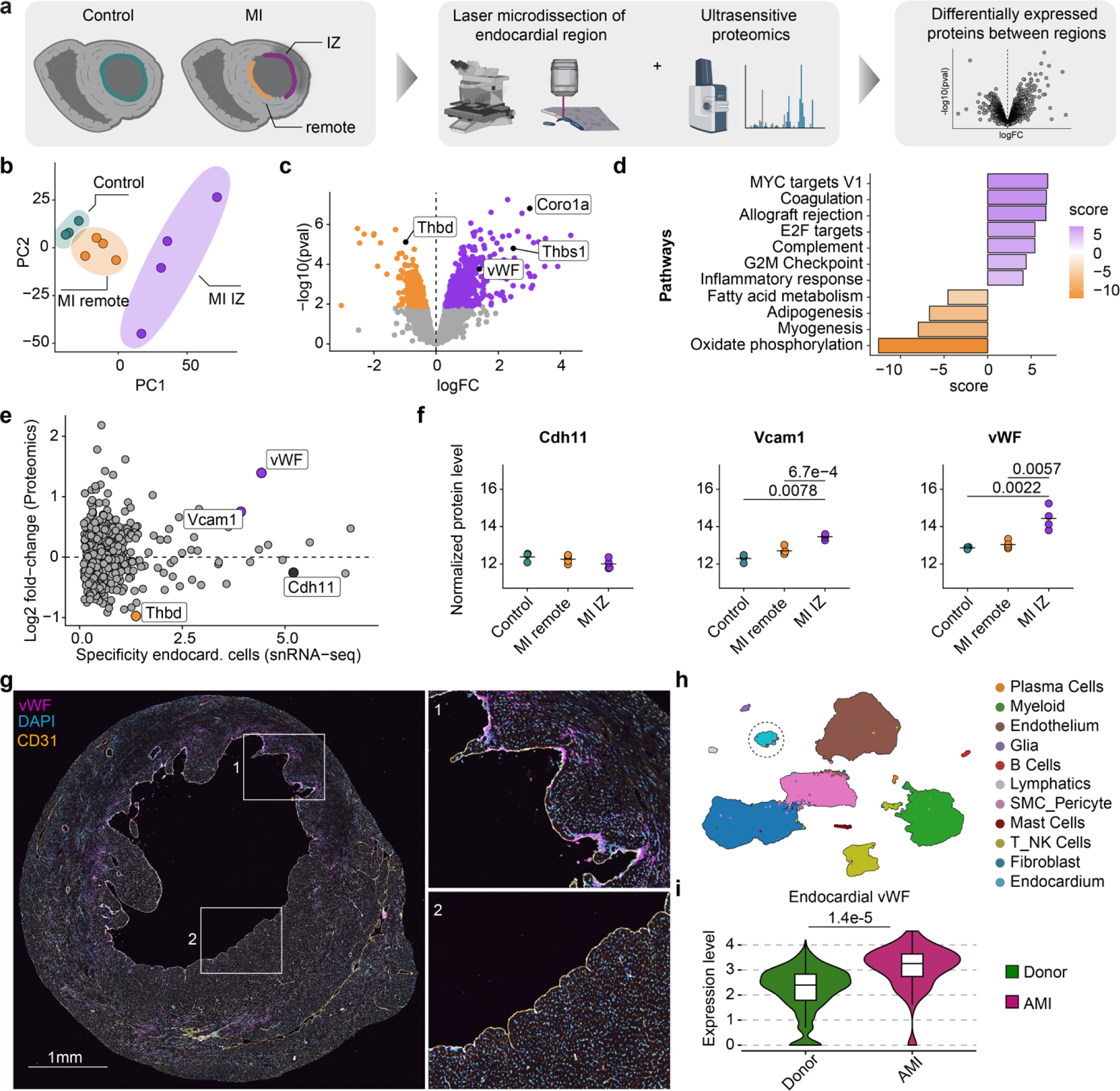
Laser capture microdissection coupled to ultrasensitive proteomics at 1 day post-MI reveals local vWF upregulation in endocardial cells. **a)** Schema for experimental design comparing endocardium of control (green), infarct zone (IZ, purple) and remote regions (orange). **b)** PCA of the three indicated experimental groups (n=3-4 biological replicates). **c)** Volcano plot of the proteomic differential comparison between infarct endocardium and remote endocardium. Significantly differentially expressed proteins are displayed in purple (upregulated in infarct endocardium) and orange (downregulated in infarct endocardium). **d)** Pathway enrichment analysis results using Hallmark gene sets of differentially expressed proteins between MI IZ and MI remote. **e)** Endocardial cell specificity analysis of differentially expressed proteins between MI IZ and MI remote. The x-axis shows specificity of gene expression based on snRNA-seq data from Calcagno et al. ^22^, while the y-axis shows log2 fold-change from differential protein expression analysis shown in c). **f)** Expression plots of three proteins of interest based on ultrasensitive proteomics. Cdh11 is a marker for endocardial cells and not differentially expressed, whereas Vcam1 and vWF show significant differential expression during acute MI. Line represents mean expression and points represent individual measurements. **g)** Representative conventional immunofluorescence staining of vWF (magenta) alongside CD31 (yellow) 24 hours after murine MI. **h)** UMAP of human cardiac cell types identified in Amrute et al. 2022. Endocardial cell cluster used for differential gene expression analysis is highlighted with a dotted circle. **i)** Violin plot of normalized RNA expression from snRNA-seq data for donor samples (n=6) and AMI samples (n=4) from Amrute et al. (2022) shows significant upregulation of vWF in human endocardial cells of AMI samples (p-value = 1.4e-05).

### Functional blocking of vWF modifies immune cell infiltration and infarct recovery

Immunofluorescence co-staining of CCR2 and vWF highlighted a strong correlation between the presence of vWF within the endocardial infarct region and locally attached or already infiltrated CCR2^+^ Mo/Mϕ after MI and myocardial ischemia/reperfusion (**Figure 6a,b, Supplementary Figure 10a-c**). Of note, immunofluorescent co-staining of platelets revealed their presence in some, but not all of these infiltration areas with expression of vWF (**Supplementary Figure 10d**). We conclude that vWF-dependent immune recruitment in this context is, at least partially, platelet-independent. To investigate the functional role of vWF in the recruitment and infiltration of myeloid cells via the endocardium during acute MI, a well-characterized polyclonal antibody (against human vWF which is also highly cross-reactive with murine vWF) was used to block vWF function in murine MI (**Figure 6c**). Control IgG or anti-vWF antibodies were injected intravenously at 0h and 24h post-MI. Blockade of vWF resulted in a significant decrease in recruited Mo/Mϕ in the infarcted myocardium as quantified by flow cytometry 2 days post-MI, whereas blood levels of Mo/Mϕ were unaltered (**Figure 6d,e**). Immunofluorescence staining demonstrated that this effect was mainly explained by dramatically reduced Mo/Mϕ within the endocardial infarct area (**Figure 6f**). Interestingly, blockade of Mo/Mϕ recruitment mediated by vWF blocking at the endocardium led to impaired healing and deteriorated long-term outcome two weeks after MI induction as shown by echocardiography (**Figure 6g-k**). Moreover, histopathological evaluation revealed more pronounced infarct thinning in anti-vWF treated mice compared to control mice (**Figure 6l-n**). Our findings using DVP and functional experiments have therefore uncovered a novel, potentially critical role of the endocardium in facilitating infiltration of myeloid cells that is likely mediated by vWF.

**Figure 6:**
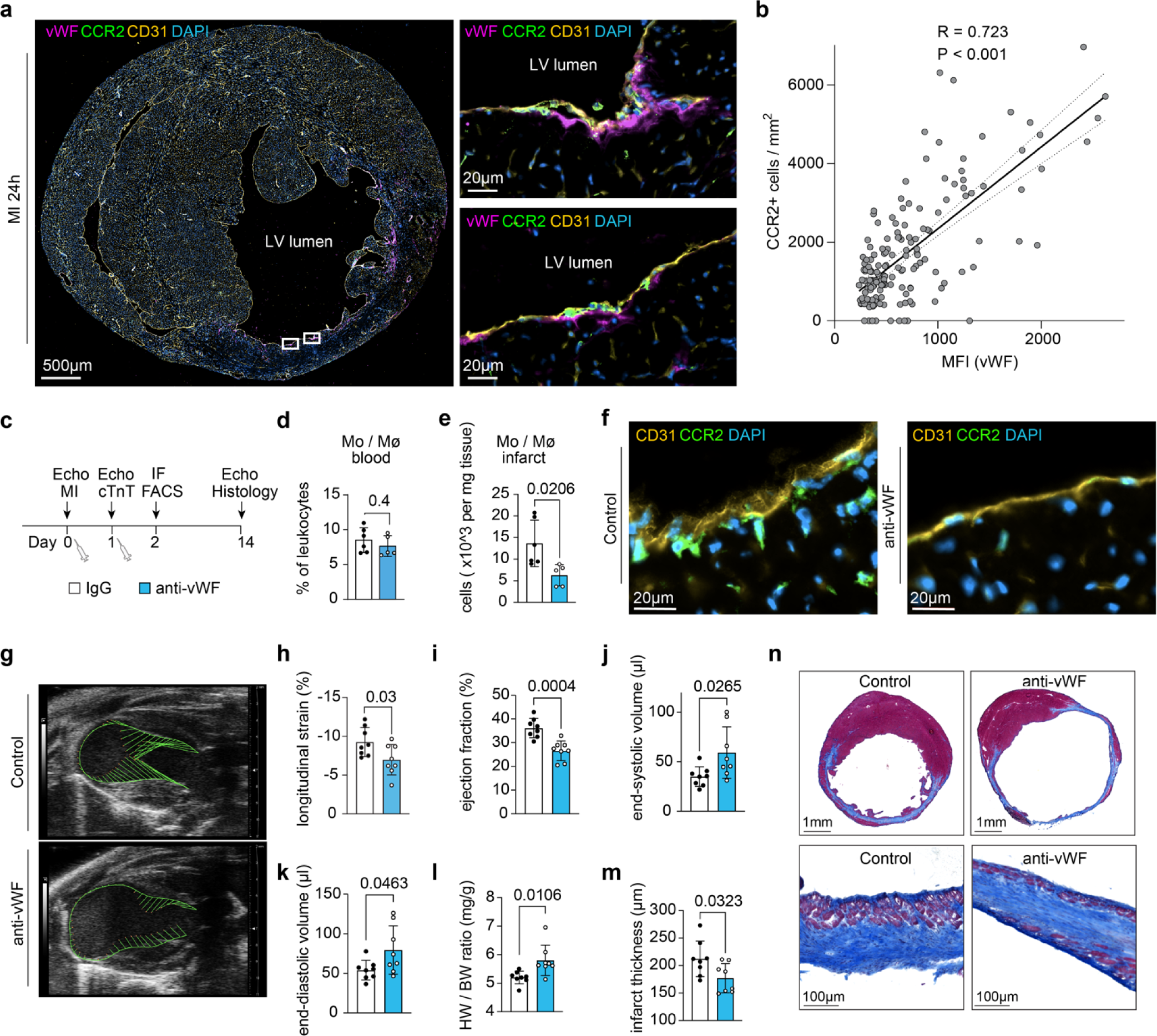
Blockade of vWF results in decreased Mo/Mϕ recruitment and impaired infarct healing. **a)** Immunofluorescence staining of vWF, CCR2, CD31 and DAPI 24h after MI. **b)** Correlation of CCR2+ cells and mean fluorescence intensity (MFI) of vWF within the endocardial infarct area. A total of 164 annotations of similar size were added across the endocardium to assess vWF MFI and the presence of CCR2+ cells within each annotation (n = 3 samples). **c)** Schema of the experimental set-up for functional vWF blocking during acute MI. **d)** Quantification of blood Mo/Mϕ 2d post-MI. **e)** Quantitative analysis of cardiac Mo/Mϕ 2d post-MI based on flow cytometry. **f)** Representative immunofluorescence stainings of CCR2+ cells (green) after both IgG and anti-vWF treatment 2d post-MI. **g)** Representative B-mode images 14d after MI induction. Green lines depict LV endocardial displacement. **h-k)** Global longitudinal strain, LV ejection fraction, end-systolic and end-diastolic volume determined by echocardiography 14d after MI induction. **l)** Heart weight to body weight ratios after organ removal 14d post-MI. **m)** Quantification of infarct thickness based on histopathological evaluation 14d after MI. Boxes in d,e and h-m represent mean±SD, points display individual measurements (n = 8 samples per group). **n)** Representative images of Masson trichrome stainings of both IgG-treated and anti-vWF treated mice 14d after infarction.

## Discussion

The heart is composed of multiple cell types which cooperate to ensure its vital function. A detailed understanding of this cellular interplay and molecular heterogeneity during homeostasis and disease is the foundation for the development of novel treatments or the improvement of existing therapies. Here, we created high resolution, transcriptomic (Molecular Cartography) and antibody-based highly-multiplexed (SeqIF) datasets and performed spatial analysis of infarct tissue in a murine MI model to explore the cellular landscape during the first four days following an infarct. While human samples can only be analyzed partially due to size restrictions and longitudinal measurements are impossible, murine models enable a comprehensive time-resolved view of the acute MI environment. This spatial multi-omics study of the acute murine MI generated a detailed spatial-temporal map of cell-type distributions within a full cross-section of the infarcted heart at subcellular resolution for the first time. Besides creating unique datasets, machine learning models and software as a foundation for future exploration, we discovered a novel mode of immune cell infiltration in a non-reperfusion MI model: We unexpectedly found endocardial attachment of CCR2+ Mo/Mϕ and recruitment into the subendocardial infarct area. To investigate the underlying mechanism, we performed ultrasensitive proteomics of endocardial cells after MI and identified vWF as a critical mediator in endocardial cells facilitating the attachment and recruitment of monocytes. This finding was further supported by functional experiments blocking vWF, which not only decreased the immune cell infiltration but also negatively impacted infarct recovery. The upregulation of vWF in endocardial cells after MI was also corroborated in human tissue. Overall, the data and findings presented in this study significantly contribute to our understanding of the inflammatory processes in the early phase of healing after MI by providing a spatiotemporal scale for cellular and molecular dynamics and will aid future studies - beyond the cardiovascular field - to investigate the diverse infiltration routes of immune cells into various organs and tissues.

## Methods

### Mouse experiments

C57BL/6NRj female mice were obtained from Janvier labs (Saint-Berthevin, France) and were studied at 10 - 12 weeks of age. Mice were housed under standard laboratory conditions with a 12h light-dark cycle and access to water and food ad libitum. All animal procedures were approved by the institutional review board of the University of Heidelberg, Germany, and the responsible government authority of Baden-Württemberg, Germany (project number G-106/19 and G-94/21).

### Minimal-invasive induction of MI

For MI induction in mice, the procedure was performed as described previously ^35^. Briefly, mice were anaesthetized with inhalation of 2 % isoflurane and placed on a Vevo imaging station connected to a Vevo 2100 system (VisualSonics, Toronto, Canada). After a brief evaluation of cardiac function, the left coronary artery was visualized. After attaching a neutral electrode, a monopolar needle controlled by a micromanipulator was inserted into the chest and placed on the coronary artery. The vessel was coagulated with high-frequency electricity using an electrosurgical station which was connected to both electrodes. After removal of the needle, successful MI was confirmed by persisting absence of a Doppler signal and akinesia in the affected part of the LV wall.

### Organ removal and preparation

Peripheral blood was collected by facial vein puncture in heparinized tubes. Hearts were excised after cervical dislocation and rinsed extensively in ice-cold PBS in order to remove remaining blood within the LV lumen and vasculature. After transverse sectioning using a scalpel, freshly dissected mouse cardiac samples were embedded in OCT compound in a plastic cassette and were immediately placed in an isopentane bath on dry ice for further processing.

### Molecular Cartography (Highly multiplexed single molecule Fluorescence In Situ Hybridization (smFISH)) of murine MI samples

10µm thick cryosections were placed within the capture areas of cold Resolve Biosciences slides. Samples were sent to Resolve Biosciences (Monheim, Germany) on dry ice for analysis. Upon arrival, mouse tissue sections were thawed and fixed with 4% v/v Formaldehyde (Sigma-Aldrich F8775) in 1x PBS for 30 min at 4 °C. After fixation, sections were washed three times in 1x PBS for one minute, followed by a one minute wash in 70% ethanol at room temperature. Fixed samples were used for Molecular Cartography™ (100-plex combinatorial single-molecule fluorescence in-situ hybridization) according to the manufacturer’s instructions and as previously described ^53^. The probes for 100 genes were designed using Resolve Biosciences’ proprietary design algorithm. **Supplementary Table 1** highlights the gene names and catalogue numbers for the specific probes designed by Resolve Biosciences.

### Image processing of Molecular Cartography data

Slides used for combinatorial single molecule FISH imaging with Molecular Cartography for 100 candidate transcripts were subsequently stained for nuclei (DAPI) and wheat-germ agglutinin (WGA), which labels plasma membranes, to facilitate cell segmentation. DAPI and WGA images as well as RNA spot tables from Molecular Cartography were processed using an in-house developed Nextflow pipeline. This pipeline is part of the nf-core community and called nf-core/molkart (https://nf-co.re/molkart/dev, revision: 81eafe9f9993d4daf16371ba3804ce9ae08053ad). Many core components of the pipeline were made available as nf-core DSL2 modules (https://nf-co.re/modules) to facilitate easy enhancement of the pipeline by other users and to enable reuse of pipeline components by Nextflow imaging pipelines in the future. The pipeline consists of multiple steps as depicted in **Supplementary Figure 3**: First, consecutive Gaussian blurring was used to fill in black grid lines from Molecular Cartography imaging using the Python tool Mindagap ^54^. Next, image stacks of DAPI and WGA were created and contrast-limited adaptive histogram equalization (CLAHE) was applied to improve contrast across stainings for automated segmentation ^55^. Training images for Ilastik Multicut (hdf5 format) and Cellpose (tiff format) were created with the use of the “create training subset” as part of the nf-core/molkart pipeline. Cell segmentation was performed using several different segmentation algorithms to compare them on cardiac images (**Supplementary Figure 4**). The segmentation algorithms used here were Deepcell Mesmer ^33^, which performs nuclear and whole-cell segmentation; Cellpose 2^34^ and Ilastik Multicut (Pixel classification + Boundary based Multicut) ^42^. For Cellpose 2, we trained a custom Cellpose model, by selecting small crops (1000 × 1000 pixels) of DAPI and WGA image stacks across the entire dataset, using a human-in-the-loop approach, as described by Pachitariu and Stringer ^34^. We used the baseline model CPx, with flow_threshold = 0.6 and cellprob_threshold = 0 to segment an initial image crop and we corrected wrong segmentations and retrained the model. This process was consecutively applied to retrain the model on new image crops until segmentation captured most cells correctly in the presented image. For Ilastik Multicut, we first trained a pixel classification model by labeling WGA signals as “membrane”, nuclear and empty cytoplasm pixels as “cell” and “background”. We selected all features and feature sizes to train the pixel classification model. Probability maps from the pixel classification were subsequently used in boundary-based segmentation with Multicut, which first uses a distance transform on the boundary probability map to create seeds for watersheds and calculate superpixels. Multicut was then performed on Superpixel edges by training a random forest edge classifier. Following segmentation, the resulting masks were size-filtered to remove extremely small and extremely large objects that don’t represent real cells. To assign spots to cells, spots are first filtered for potential duplicates using the Mindagap duplicatefinder function, which filters potential duplicate RNA spot calls along black grid lines. Deduplicated RNA spots are then assigned to segmentation masks using spot2cell, which creates cell-by-feature tables containing counts for each transcript per cell, and cell shape properties. Finally, quality control metrics of all relevant steps are collected and compiled for inspection via MultiQC. Images and spots were visualized using the Napari toolkit, which enables fast and interactive plotting of large imaging data ^56^.

### Single-cell analysis of Molecular Cartography data

Cell-by-feature matrices from nf-core/molkart were imported into R and processed using the Seurat package (version Seurat_5.0.1) ^57–60^. We filtered out cells with less than 20 and more than 4000 RNA counts. We also filtered outlier cells based on their segmentation mask shape with extent < 0.25 and solidity <0.75 (as estimated by the regionprops_table function from scikit-image package in Python) to ensure only high-quality segmented cells are included in the analysis. Cell transcript profile counts were normalized using SCtransform in Seurat ^58,60–62^, principal components were calculated and the first 30 principal components were used for integration of samples across time using the IntegrateLayers function in Seurat with the method set to “HarmonyIntegration” ^63^. Harmony embeddings were then used for uniform manifold approximation (UMAP) and cluster identification using shared nearest-neighbour analysis using the first 30 harmony dimensions. We used cell type label transfer to annotate the Molecular Cartography cells using a reprocessed snRNA-seq reference from Calcagno et al. 2020 ^22^. To reprocess the snRNA-seq data from Calcagno et al., we performed Seurat analysis on the raw data as described by the authors in the original publication. The transferred labels from Calcagno et al. were used to guide manual annotation of cells into cell types and states and produce final labels for all cell clusters. To improve the annotation of endocardial cells, we additionally manually labeled the endocardial region in all DAPI + WGA images using QuPath ^64^. We exported endocardial masks as GeoJSON files in QuPath and processed them using the sf package in R to overlay centroid positions of cell masks. Cells that overlapped the endocardial region and expressed Pecam1 (normalized count > 0) or were clustered into an Npr3 positive cell cluster were considered as endocardial cells.

We characterized structural patterns in immediate cellular neighborhoods by extracting cell-type to cell-type relationships with MISTy (R package mistyR, version 1.99.9) ^43^. MISTy is a multiview framework for analysis of spatial omics data by identification of robust relationships within the data coming from different spatial contexts. Based on Molecular Cartography assigned cell types, we represent each cell-type intrinsically as a one-hot encoded vector. To capture the structure of the spatial neighborhood of each cell, we added a paraview with a radius of 125µm. The paraview captures the neighborhood composition by distance-weighted sum of one-hot encoded representation of the cell types in the surrounding of each cell. The weights are calculated by a radial basis function with parameter equation to the chosen radius. Subsequently, a MISTy model was trained using the same view composition for each sample independently. The MISTy models are trained on the task of predicting each intrinsic cell-type by using all variables from the paraview. The MISTy output consists of the amount of variance explained per target and importances of each predictor-target interaction. The importance of each interaction was standardised to 0 mean and unit standard deviation across all predictors for a given target. The performance and interaction results were aggregated per time point, filtered to exclude all targets with variance explained less than 5% and relationships with importances lower than 0.4. MISTy captures robust relationships on a global scale, i.e., consistent across the whole slide. Additionally MISTy can learn not only simple linear relationships, but also complex non-linear relationships. To linearly approximate the sign of the remaining relationships and estimate their consistency across each slide, we calculated the correlation between the predictor variables from the paraview and the target variables from the intraview. While strong correlations are indicative of linear and consistent relationships, correlations close to 0 point towards non-linearity or heterogeneity of the form of the interaction across the slide, warranting a more targeted local bivariate spatial analysis. We used Liana+ (version 1.0.4) to calculate spatially-informed local bi-variate metrics between pairs of cell types of interest. Similar to MISTy, we used one-hot encoded cell-type vectors and calculated the cell neighbourhoods using a Gaussian kernel with a cutoff of 0.1 and a bandwidth of 125µm and l1 standardization of terms. We then use the lr_bivar function in Liana+ to calculate the normalized weighted product between two cell-type vectors as input. Interactions were visualized by plotting local scores on tissue coordinates. To calculate Euclidean distances between pairs of cells, we used the Scipy spatial packages cdist function ^65^. Statistical testing on distances was performed with ANOVA with a linear OLS Model and post hoc t-tests with Bonferroni correction using the statsmodel API in Python ^66^.

### Sequential Immunofluorescence (SeqIF) imaging using Lunaphore COMET™ platform

For SeqIF stainings, samples were sectioned on a cryotome (8μm) and collected on adhesion slides (Epredia™ SuperFrost Ultra Plus™ GOLD, Fisher Scientific) and dried on a 37°C heat plate for 15 min. After storage at −80°C, sections were brought to room temperature and were incubated in 4% formaldehyde for 40 min at room temperature. Samples were washed for 5 min at room temperature in Multistaining buffer (BU06, Lunaphore Technologies) followed by incubation with Multistaining Buffer supplemented with 0.2% Triton for 20 min at room temperature. Subsequently, slides were stored in Multistaining Buffer till use. Slides were dried off and placed into COMET™ stainers with microfluidic chips positioned on top of the tissue section as described by Rivest et al. ^48^. Antibody mixes were prepared by diluting the stock antibody solutions in Intercept-T20 to help with blocking of non-specific binding (**Supplementary Table 2**). Automated Sequential Immunofluorescence (SeqIF™) staining and imaging were performed on the COMET™ platform (Lunaphore Technologies). Slides underwent 13 cycles of iterative staining and imaging, followed by elution of the primary and secondary antibodies ^67^. The 16-plex protocol template was generated using the COMET™ Control Software and reagents were loaded onto the device to perform the SeqIF protocol. A list of primary antibodies with corresponding incubation times can be found in **Supplementary Table 2.** Secondary antibodies were used as a mix of 2 species-complementary antibodies: Alexa Fluor™ Plus 647 goat anti-rabbit (Thermo Scientific, cat no: A32733, 1/250 dilution) and Alexa Fluor™ Plus 555 goat anti-rat (Thermo Scientific, cat no: A48263, 1/250 dilution). Nuclear signal was detected using DAPI (Thermo Scientific, cat no: D1306, 1/1000 dilution) by dynamic incubation of 2 min. All reagents were diluted in Multistaining Buffer (BU06, Lunaphore Technologies). The elution step lasted 2 min for each cycle and was performed with Elution Buffer (BU07-L, Lunaphore Technologies) at 37°C. The quenching step lasted for 30s and was performed with Quenching Buffer (BU08-L, Lunaphore Technologies). The imaging step was performed with Imaging Buffer (BU09, Lunaphore Technologies). The output from the COMET™ platform is a stitched and registered, multi-stack ome.tiff file, that was directly used for further processing with MCMICRO ^49^ and downstream applications.

### Image processing and analysis of SeqIF (Lunaphore COMET™) data

Post-acquisition registration of full slide images was needed for 2 images due to interrupted runs and was performed with Palom (https://github.com/labsyspharm/palom). Processing of multi-stack ome.tiff files was performed using a modified version of MCMICRO ^49^. Autofluorescence signal subtraction was performed on a pixel-level basis using the following formula: 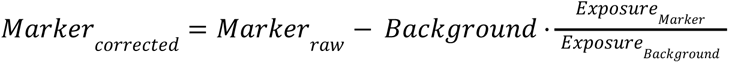. Preprocessing of the images to improve segmentation was performed using contrast-limited adaptive histogram equalization (CLAHE) on the DAPI and membrane (WGA, wheat-germ agglutinin) channels. Cell segmentation was performed similar to Molecular Cartography data using CLAHE-adjusted images to train a custom Cellpose 2 model via the human-in-the-loop approach. Feature quantification was performed on the autofluorescence-subtracted image based on the labeled segmentation masks. To assign cell phenotypes, we utilized a pixel-level clustering workflow using self-organizing maps (SOMs) implemented via the Pixie pipeline ^50^. Therefore, we manually annotated the image region containing the heart using QuPath (0.4.3, MacOS version) ^64^ and set all background pixels to 0 across all channels to reduce the number of pixels to be processed by Pixie. We then performed pixel clustering with 10 functional markers: Ankrd1, aSMA, CCR2, CD31, CD45, CD68, Mpo, Pdgfra, Tnnt2 and Trem2 across all 9 COMET images using 5% of pixels subsets to train the SOM. Pixel meta clusters were visualized using the Pixie Jupyter widget and 100 SOM clusters were manually merged and visualized as pixel phenotype maps for validation. Pixel clusters were subsequently used in a second clustering step with Cellpose masks to assign cell phenotypes via SOM clustering across all images. Similar to pixel clustering, 100 SOM clusters were manually merged into cell phenotypes and each cell segmentation mask was assigned a phenotype. To quantify cell-type abundances in different regions of the heart (endocardial layer, epicardial layer, infarct core, border zones), we manually annotated regions in QuPath and used them to subset cells based on their presence or absence in the annotation. For the quantification of myeloid cells in the endocardium, we included cells in the LV lumen, only if they were in direct contact with endocardial cells.

### Laser-microdissection coupled to ultrasensitive proteomics

To collect cells with the Leica Laser Microdissection 7 (LMD7) microscope three reference points are required to triangulate the shape coordinates into laser cutting coordinates. These reference points were etched using the LMD7 on the membrane of the Leica Frame slides (Order # 11600294) before the tissues were placed on them. These etchings were easily recognizable features to which we could go back and designate them as reference points. Five micrometer thick tissue sections from fresh frozen heart tissue were prepared similarly to the SeqIF samples and were placed onto Leica Frame slides with reference points. Slides were stained using DAPI, WGA and CD31 (as described in the IF section) and imaged using a Zeiss Axioscan7. Stitched images of whole hearts were imported and annotated in QuPath for downstream laser capture microdissection. Endocardial regions to be collected by the LMD7 were annotated using the QuPath’s brush tool with a brush diameter of approximately 20µm, centered around endocardial cells with 10µm to each side as buffer for the laser cutting. For control hearts, only one endocardial group was annotated, while for hearts 1-day post-infarct, endocardial cells in the infarct region and endocardial cells in the remote region were labelled separately. QuPath annotations were then exported as GeoJSON files, which were further processed using the Qupath_to_LMD scripts (https://github.com/CosciaLab/Qupath_to_LMD). The script assigns annotation classes to wells of the 384-well plate. It uses the py-lmd package from Sophia Madler and colleagues (https://github.com/MannLabs/py-lmd) to transfer GeoJSON polygons into LMD readable .xml files.

### Laser capture microdissection

We used the Leica LMD 7 system and Leica Laser Microdissection V 8.3.0.08259 software to collect tissue contours. Tissue was cut with a 63x objective (HC PL FLUOTAR L 63x/0.70 CORR XT) in brightfield mode. Laser settings were: Power 60; Aperture 1; Speed 25; Middlepulse Count 2; Final Pulse 5; Head Current 39-41%; Pulse Frequency 2028, and offset of 105. Contours were cut and sorted into a low-binding 384-well plate (Eppendorf 0030129547) configured over the ‘universal holder’ function.

### Sample preparation for LC-MS analysis

To collect tissue pieces stuck on the sides of the 384 wells, we washed them down with 30µl of acetonitrile, briefly vortexed and vacuum dried (15 min at 60°C). We added 2µL of Lysis Buffer (0.1% DDM, 5mM TCEP, 20mM CAA resuspended in 100mM TEAB; pH 8.5) to each well, closed the plate with a PCR ComfortLid (Hamilton), and heated at 95°C for 60 min. We added 1µl of LysC (2 ng/µl in water), and incubated for at least 2 hours at 37°C. Subsequently, 1µl of trypsin was added (1 ng/µl in water) and the samples were incubated overnight at 37°C. The next day the samples were vacuum dried before peptide clean-up. Peptide clean-up took place with Evotips (Evosep, Odense, Denmark) following manufacturer’s recommendations. Briefly, the Evotips (EV2013, Evotip Pure, Evosep) were washed with Buffer B (99.9% ACN, 0.1% FA) and then Buffer A (99.9% water, 0.1% FA); then activated with isopropanol. Digested tissue samples were resuspended in Buffer A, loaded into the tips, washed with Buffer A once, and then eluted with Buffer B into a 96-well plate (Thermo Fisher Scientific, AB1300), and vacuum dried. Samples were stored at −20°C until liquid chromatography-mass spectrometry (LC-MS) analysis. For LC-MS analysis, 4.2µl of MS loading buffer (3% acetonitrile, 0.1% TFA in water) was added, from which 4.0µl were finally injected into the mass spectrometer.

### Liquid chromatography-mass spectrometry (LC-MS) analysis

LC-MS analysis was performed with an EASYnLC-1200 system (Thermo Fisher Scientific) connected to a trapped ion mobility spectrometry quadruple time-of-flight mass spectrometer (timsTOF SCP, Bruker Daltonik) with a nano-electrospray ion source (CaptiveSpray, Bruker Daltonik). Peptides were loaded on a 20cm home-packed HPLC column (75µm inner diameter packed with 1.9µm ReproSil-Pur C18-AQ silica beads, Dr. Maisch). Peptides were separated using a linear gradient of 21 min and analysed in dia-PASEF mode.

### Proteomics data analysis

We used DIA-NN (1.8.2) for dia-PASEF raw file analysis and the generated libraries were used for mouse proteins (UNIPROT mouse released in 2021) and known contaminants ^68^. Deep learning-based spectra, RTs and IMs prediction were enabled for the appropriate mass range of 300-1200 m/z. N-terminal M excision and cysteine carbamidomethylation were enabled as fixed modifications. A maximum of 2 miscleavages were allowed, and the precursor charge was set to 2-4. DIA-NN was operated in the default mode with minor adjustments. Briefly, MS1 and MS2 accuracies were set to 15.0, scan windows to 0 (assignment by DIA-NN), isotopologues were enabled, MBR, heuristic protein inference and no shared spectra. Proteins were inferred from genes, neural network classifiers were set to single-pass mode, quantification strategy as ‘Robust LC (high precision)’. Cross-run normalization was set to ‘RT-dependent’, library generation as ‘smart profiling’, speed and RAM usage as ‘optimal results’. Protein lists were filtered for missing values by group, requiring at least two observed values in the control group or 3 observed values in the MI_remote or MI_IZ group. We also filtered out known contaminants based on previously described contaminants libraries ^68^. Differential protein expression analysis was performed on data normalized using variance stabilizing normalization using empirical Bayes statistics in limma ^69^. Proteins with an FDR lower than 0.05 were considered as significantly differentially expressed. Overlap in the proteins differentially expressed between conditions was visualized using Upset plots with the ComplexUpset package ^70,71^. Endocardial specificity of differentially expressed proteins was compared by correlating log-fold changes from proteomics results with log-fold changes from marker gene estimations (Seurat’s FindMarker function) for endocardial cells at day 0 from the reprocessed Calcagno et al. dataset.

### Echocardiography

Echocardiographic analyses were performed in conscious mice using a Vevo 2100 ultrasound system (VisualSonics). LV end-diastolic volume, end-systolic volume, and ejection fraction were measured based on the left parasternal long-axis view and were acquired using VevoLab software (VisualSonics). Global longitudinal strain was quantified in the longitudinal axis by speckle tracking using VevoStrain software (VisualSonics). Investigators were blinded to the sample group allocation during the experiments and analyses.

### Flow cytometry

Single-cell suspensions of infarcted hearts were obtained by mincing the tissue with fine scissors and digesting it with a solution containing 450 U/mL collagenase I, 125 U/mL collagenase XI, 60 U/mL DNase I, and 60 U/mL hyaluronidase (MilliporeSigma) for 1 hour at 37°C while shaking. For flow cytometry of blood samples, erythrocytes were lysed in RBC lysis buffer (Miltenyi Biotec). The fluorescent antibodies are described in **Supplementary Table 3**. Flow cytometry was performed on a FACSVerse (BD Biosciences). Data was analyzed using FlowJo software. Mo/Mϕ were identified as CD45+, Lin-(CD19;CD4;NK1.1;Ly6G;Ter119), CD11b+.

### Histology

Histopathological evaluation of LV remodelling was performed on day 14 after MI induction. Hearts were excised and rinsed in PBS. After transverse sectioning using a scalpel, hearts were then embedded in Tissue-Tek OCT compound (Sakura) and placed in 2-methylbutane (Honeywell) on dry ice. Hearts were stored overnight at −80°C and sectioned using a cryostat (9μm thickness). Tissue sections were stained with the Masson’s Trichrome Stain Kit (MilliporeSigma) according to the manufacturer’s instructions. Scar thickness was averaged from 5 measurements in short axes in a blinded fashion.

### Conventional immunofluorescence stainings

For conventional immunofluorescence staining, samples were sectioned on a cryotome (8μm), collected on adhesion slides (Epredia™ SuperFrost Ultra Plus™ GOLD, Fisher Scientific) and dried on a 37°C hot plate for 15 min. After storage at −80°C, sections were brought to room temperature and incubated in 4% formaldehyde at room temperature for 40min. Sections were permeabilized for 20min, blocked with 5% BSA for 1h, and stained overnight with primary antibodies for CD31, vWF, CCR2, CD68 or CD41 in 1% BSA staining buffer. On the following day, sections were washed and stained for 1h with the corresponding secondary antibodies combined with the labeled antibody against WGA. After washing, sections were stained with 300nM DAPI (Thermo Fisher Scientific, D1306) for 10min, washed again, and covered with a mounting medium. For simultaneous staining of CCR2 and vWF, sections were fixed, permeabilized and blocked as described above. After incubating primary antibodies for CD31 and CCR2 overnight, vWF antibody was labeled using a secondary antibody labeling kit (FlexAble CoraLite® Plus 750 Antibody Labeling Kit for Rabbit IgG, Proteintech). Labeling was carried out in accordance with the manufacturer’s instructions. In brief, vWF primary antibody was incubated with FlexLinker and FlexBuffer for 5min. FlexQuencher was added and incubated for 5 additional minutes. 1% BSA staining buffer was added and tissue sections were incubated with the labeled primary antibody. On the next day, sections were washed and stained with WGA conjugated to Alexa Fluor 488 (Thermo Fisher Scientific, W11261) for 1h. Following the WGA incubation time, sections were stained with 300nM DAPI (Thermo Fisher Scientific, D1306) for a duration of 10min, washed again and covered with a mounting medium. Images were captured using an Axio Observer (Zeiss) fluorescence microscope and analyzed using QuPath (0.4.3, Windows version). An overview of antibodies used for conventional immunofluorescence stainings can be found in **Supplementary Table 4**.

## Data availability

All relevant images and data for Molecular Cartography, SeqIF and DVP described in this study are publicly available via Synapse (project SynID : syn51449054): https://www.synapse.org/#!Synapse:syn54235747

## Code availability

All code to process data and produce the results presented in this manuscript is available on Github: https://github.com/SchapiroLabor/mi_spatialomics.

## Acknowledgements

The authors gratefully acknowledge the data storage service SDS@hd supported by the Ministry of Science, Research and the Arts Baden-Württemberg (MWK), support by the state of Baden-Württemberg through bwHPC and the German Research Foundation (DFG) through grant INST 35/1314-1 FUGG, INST 35/1503-1 FUGG and INST 35/1597-1 FUGG. F. W. was supported by a Walter-Benjamin grant from the Deutsche Forschungsgemeinschaft (DFG 516668179) and the German Federal Ministry of Education and Research (BMBF 01ZZ2004). J.N. and F.C. acknowledge support by the Federal Ministry of Education and Research (BMBF), as part of the National Research Initiatives for Mass Spectrometry in Systems Medicine, under grant agreement no. 161L0222. D.S. is supported by the German Federal Ministry of Education and Research (BMBF 01ZZ2004). We thank Wouter-Michiel Vierdag for his help with Napari, Daniel Dimitrov for help with running Liana+ and Aurélien Dugourd for help with differential protein expression analysis. F.S., J.A., K.J.L. and F.L. acknowledge support by the Leducq foundation (ImmunoFib-HF). F.S., C.H. and F.L. are supported by the DFG Collaborative Research Centre 1531 (subproject B07) and DFG Graduate Research Training Group 2727. N.H. is supported by the MD/PhD program from the Faculty of Medicine, Heidelberg (MFHD). F.L. is thankful for support from the DFG Heisenberg Programme. J.T. is supported by the “Bruno and Helene Jöster Stiftung”. This work is also supported by the Health + Life Science Alliance Heidelberg Mannheim and received state funds approved by the State Parliament of Baden-Württemberg.

## Author contributions

F.W., F.S., F.L. and D.S. designed the study. F.S. performed mouse experiments and tissue embedding/sectioning. F.S., T.T., and N.H. performed conventional immunofluorescence stainings. F.S. and C.H. performed flow cytometry experiments. F.W., K.B., F.S., and M.A.I.A. performed data analysis of molecular cartography data. J.T provided guidance for neighbourhood analysis and performed MISTy analysis under supervision of J.S.R.. F.W., K.B., F.S. and T.T. performed SeqIF experiments and data analysis. J.N. performed ultrasensitive proteomics experiments. Proteomics data analysis was done by J.N., F.W., F.S. and M.N. under supervision of F.C.. J.A. and K.J.L. provided and analyzed snRNA-seq data from human AMI samples. F.S. performed functional blocking experiments for vWF. F.W., F.S., N.F., F.L., and D.S. wrote the manuscript with input from all authors.

## Supplementary Tables

**Supplementary Table 1: Transcript panel design for Molecular Cartography.**

Overview of the custom 100 gene panel designed for Molecular Cartography. Genes were selected based on a combination of expert knowledge, evidence from single-cell studies and functional relevance. The column “Function” describes the main reason for inclusion: EMT = Endothelial-to-mesenchymal transition, cell_type_marker = gene is specifically expressed in scRNA-seq data, MI_pathway = prior evidence in myocardial infarction studies, apoptosis; inflammation; WNT_signaling = protein is involved in these pathways. “Reference_id” and “url” point towards studies highlighting the relevant gene.

**Supplementary Table 2: Antibody panel information for Lunaphore COMET, conventional IF and FACS experiments.**

Complete list of all primary and secondary antibodies used for sequential immunofluorescence experiments in this study. “Cycle” refers to the imaging cycle the antibody was used in, “Host” is the species the antibody was raised in. Cd11b* = This antibody was used in the imaging but did not pass quality control and was therefore not used in any analysis in this study.

**Supplementary Table 3: Primary and secondary antibodies used in conventional immunofluorescence experiments.**

Complete list of all primary and secondary antibodies used for conventional immunofluorescence experiments in this study.

**Supplemental Table 4: Antibodies used in flow cytometry experiments.**

Complete list of all antibodies used in flow cytometry experiments in this study.

