## Supplementary Figures for "Spatial omics of acute myocardial infarction reveals a novel mode of immune cell infiltration"

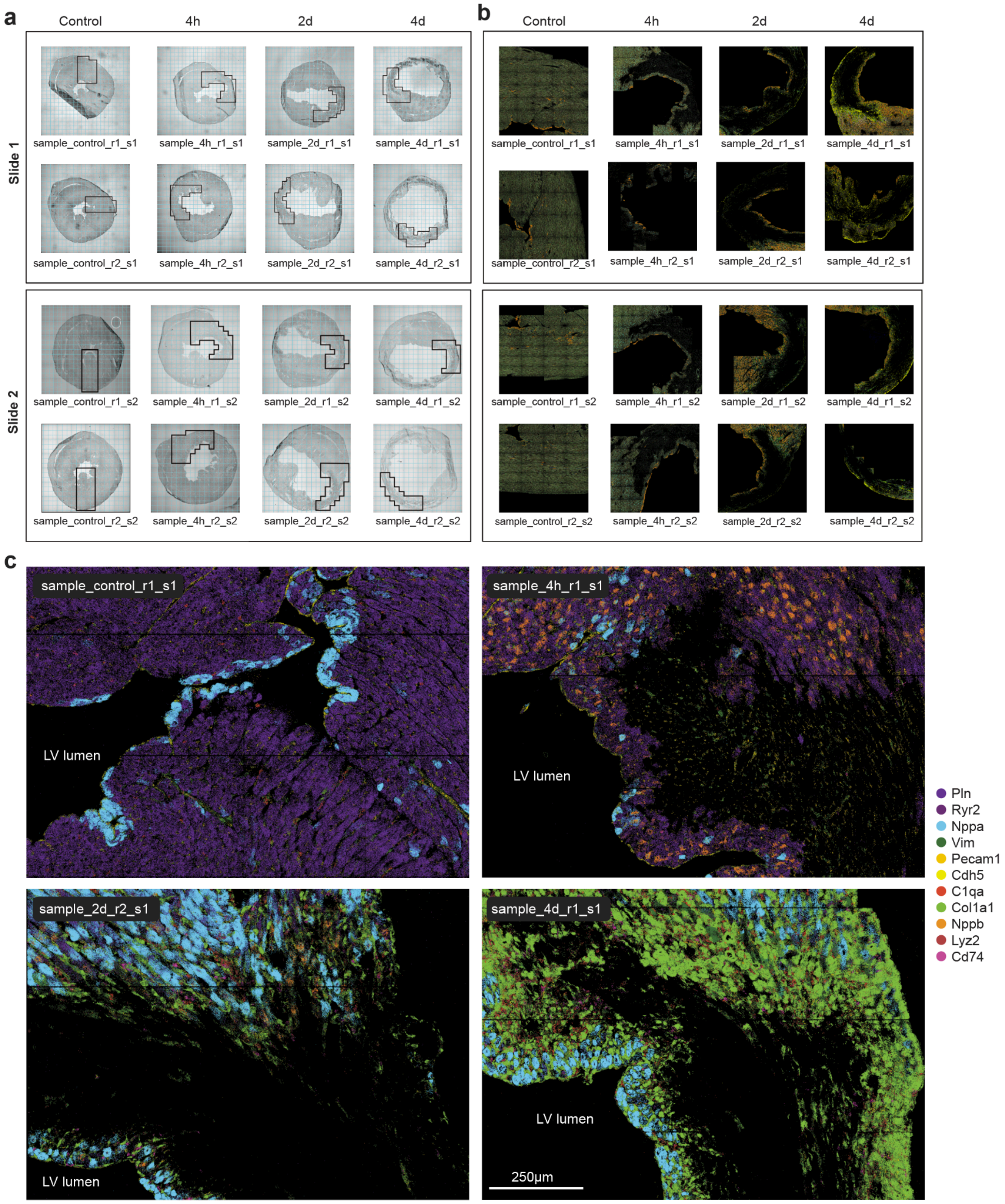


**Supplementary Figure 1: Region of interest (ROI) selection and spot distribution of mouse heart sections from Molecular Cartography. a)** Brightfield images of transverse sections of mouse hearts at different time points during acute MI (Control = prior to infarct). Black rectangles highlight regions selected for Molecular Cartography. **b)** Molecular Cartography RNA spots (100-plex) for corresponding regions highlighted in a). Regions with low spot density within the tissue at 4h, 2d and 4d demarcate infarct regions with cell death, apoptosis and RNA degradation. Note that images in b) show scatterplots of RNA spot centroid positions after spot calling by Resolve Bioscience and not the raw FISH signals. **c)** Exemplar regions of RNA expression maps of indicated markers over all 4 timepoints.

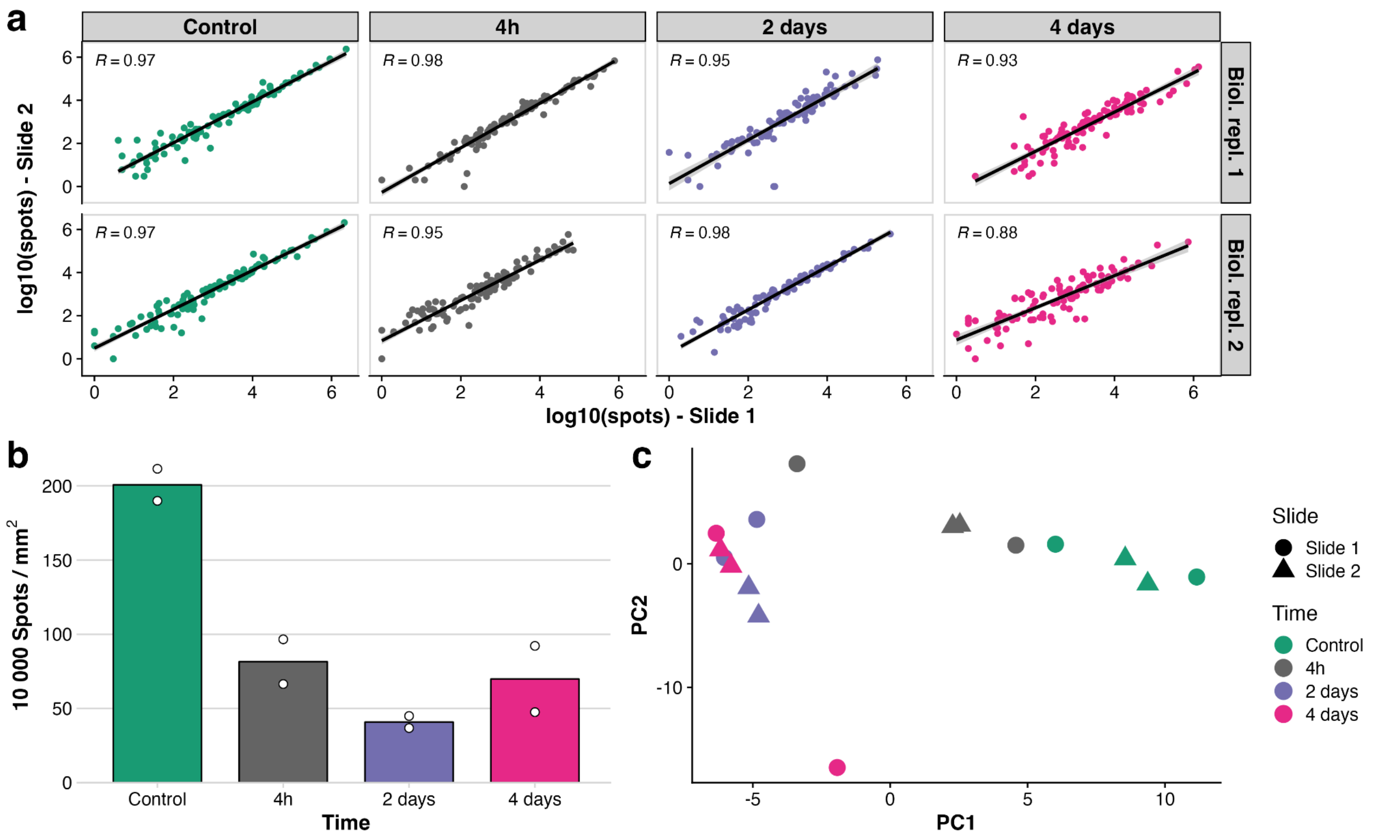


**Supplementary Figure 2: Quality control of Molecular Cartography RNA spot count data. a)** Correlation between technical replicates across two runs of Molecular Cartography across four timepoints of acute MI. R: Pearson correlation coefficient of log10 spot sum between technical replicates. **b)** Relative number of RNA spots identified per square micrometre of tissue measured using Molecular Cartography. **c)** Principal component analysis of samples by total RNA spots per gene. The outlier samples for the 4-hour and 4-day time points both have multiple small image tiles with spots missing due to failing quality control during spot calling by Resolve Bioscience.

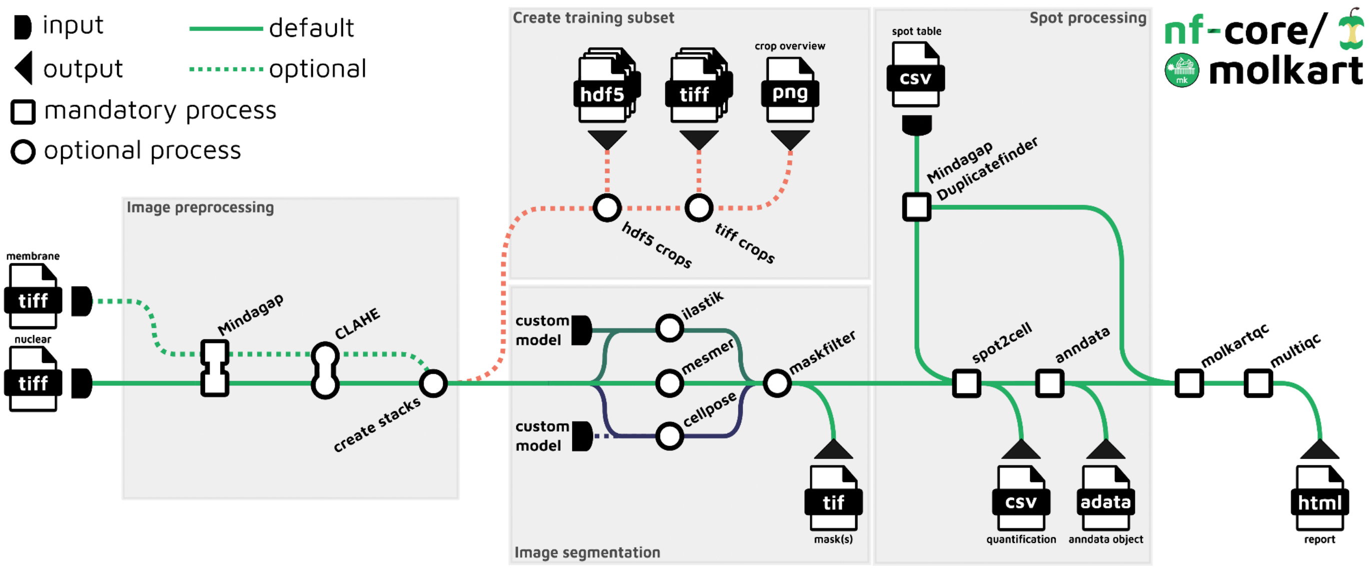


**Supplementary Figure 3: Metro map for nf-core/molkart pipeline for processing Molecular Cartography data using Nextflow.** Image data is preprocessed using Mindagap to fill in black gridlines between tiles and Contrast-Limited Adaptive Histogram Equalization (CLAHE) to improve contrast of nuclei and membrane stains across tiles. Preprocessed nuclear and membrane stains are then stacked into one image file and can be used to create image crops for training machine learning cell segmentation models or directly used for cell segmentation with either one of three segmentation methods (ilastik, Mesmer, Cellpose). The table of RNA spots is processed with Mindagap Duplicatefinder to remove potential duplicate spots near tile borders. Deduplicated spots are quality controlled using molkartqc and Multiqc and assigned to cell masks using spot2cell. The resulting cell-by-gene table is converted to an anndata object and returned to the user for downstream processing.

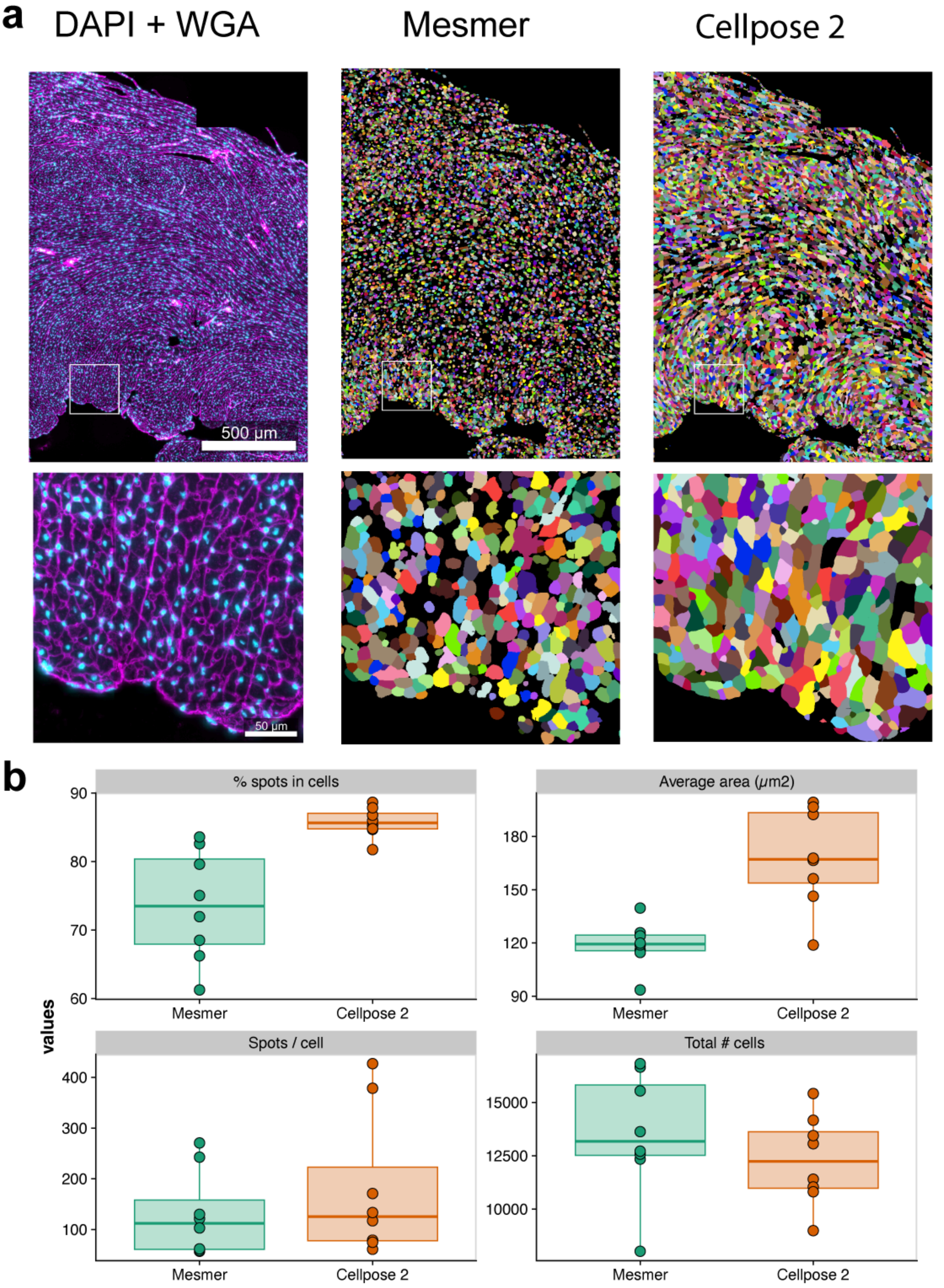


**Supplementary Figure 4: Comparison of segmentation methods for Molecular Cartography heart sections. a)** Immunofluorescence images for nuclei (DAPI) and cell membranes (WGA) on a control mouse heart section from Molecular Cartography alongside segmentation results using two different segmentation strategies are shown. **b)** Molecular Cartography cell statistics across the two different segmentation strategies.

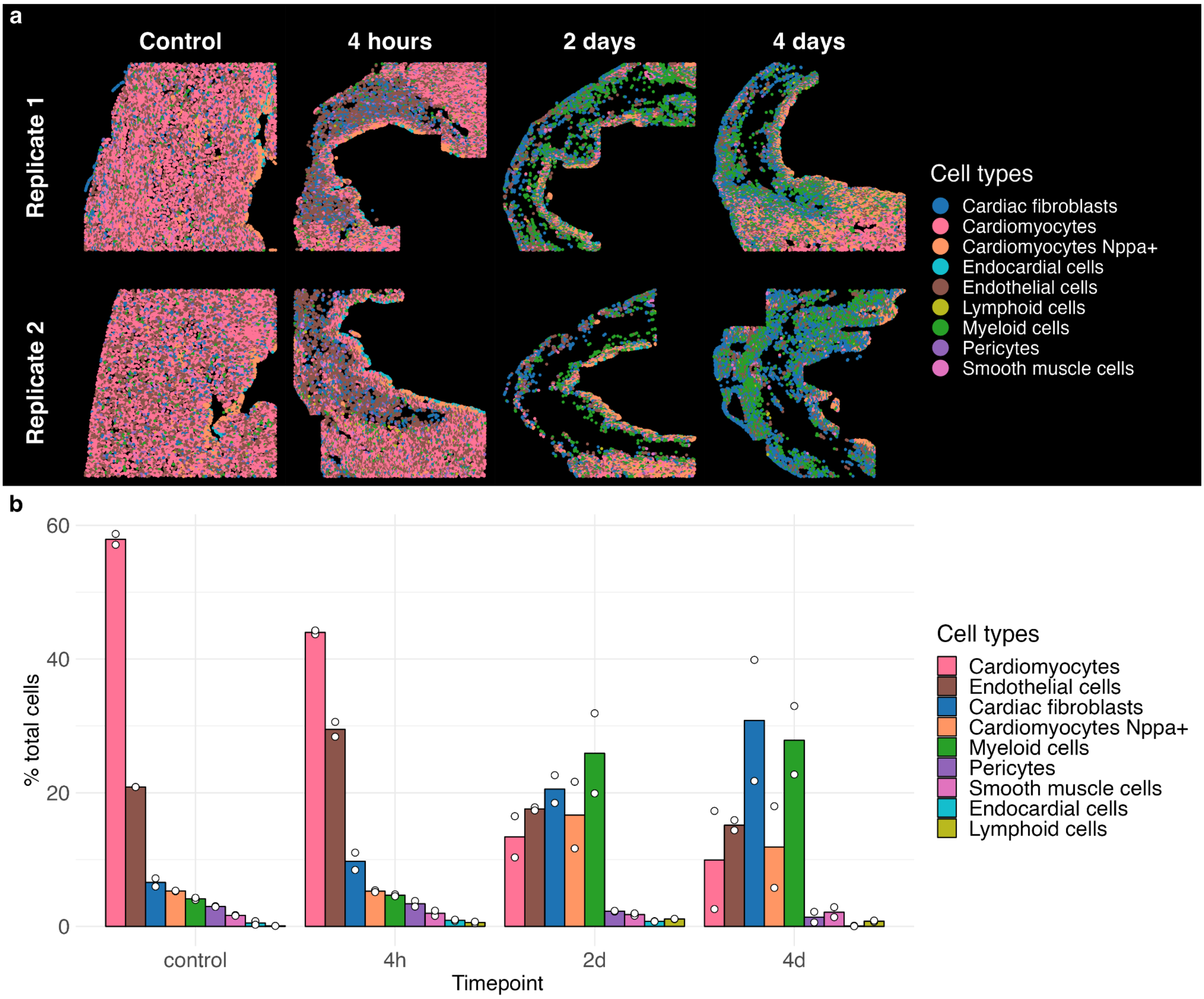


**Supplementary Figure 5: Spatial cell type distribution and composition changes in Molecular Cartography samples during acute MI. a)** Spatial distribution of cell-types in Molecular Cartography samples at four points each with two biological replicates. **b)** Cell-type composition across acute myocardial infarction as quantified by Molecular Cartography. Barplots show mean percentage, points represent individual replicate measurements.

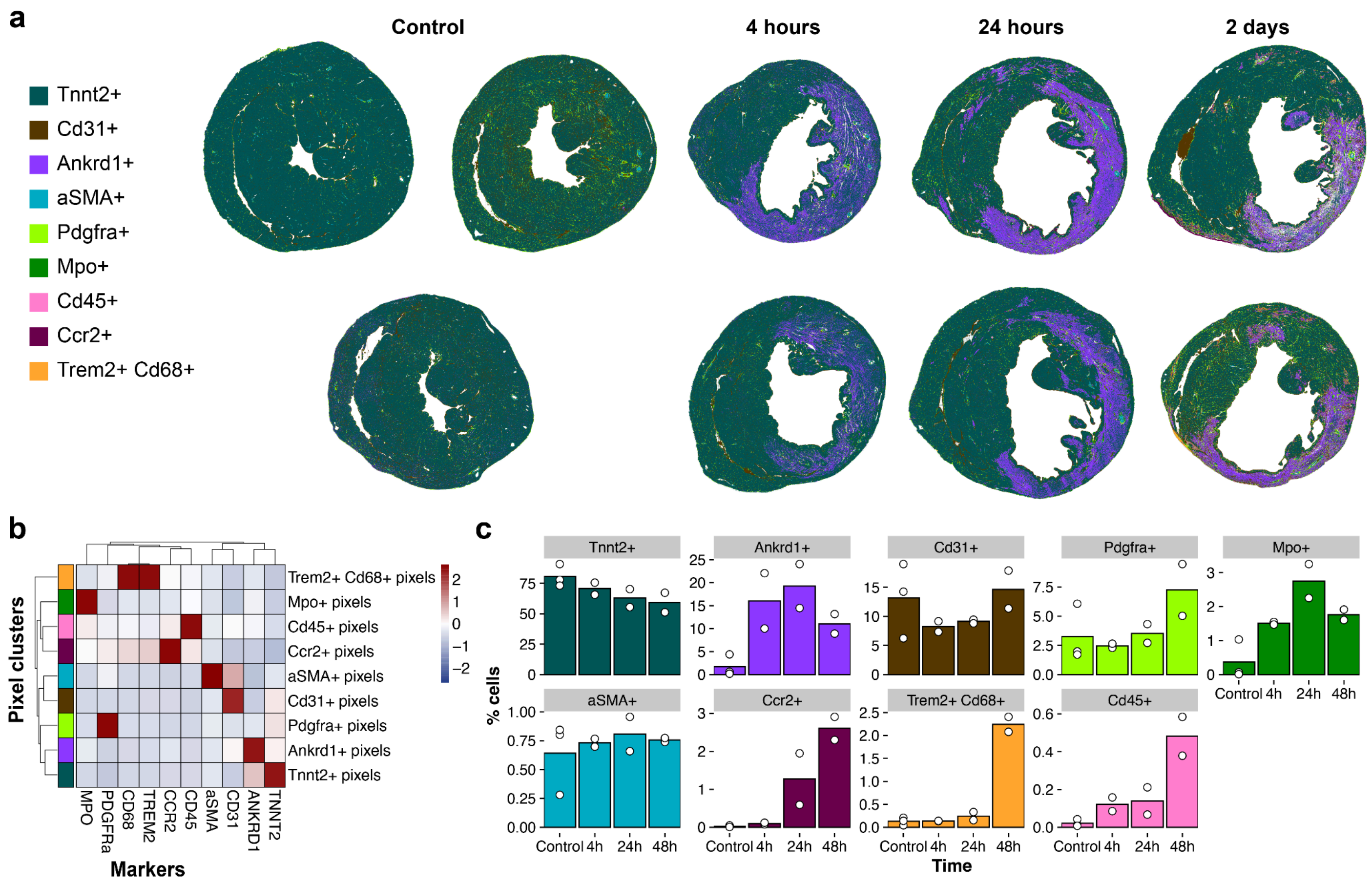


**Supplementary Figure 6: SeqIF pixel clusters and cell phenotypes across acute MI time series. a)** Pixel phenotype map for mouse heart images produced with SeqIF during a time course of acute myocardial infarction. Pixels were clustered using self-organizing maps leveraging Pixie and colored according to their assigned pixel cluster. A total of 9 different pixel clusters were classified. **b)** Heatmap for quantified marker expression in pixel phenotype clusters. Colors for pixel clusters correspond to visualization in a. **c)** Quantification of pixel phenotypes across acute MI reveals strong reduction in Tnnt2+ pixels, increase in Ankrd1+ pixels and an increase in pixel clusters for myeloid cells (CD45+, Mpo+, CCR2+, Trem2+, CD68+) during the first four days of MI. Colors correspond to pixel phenotypes visualization in a. Bars represent mean values from two biological replicates and points represent individual measurements.

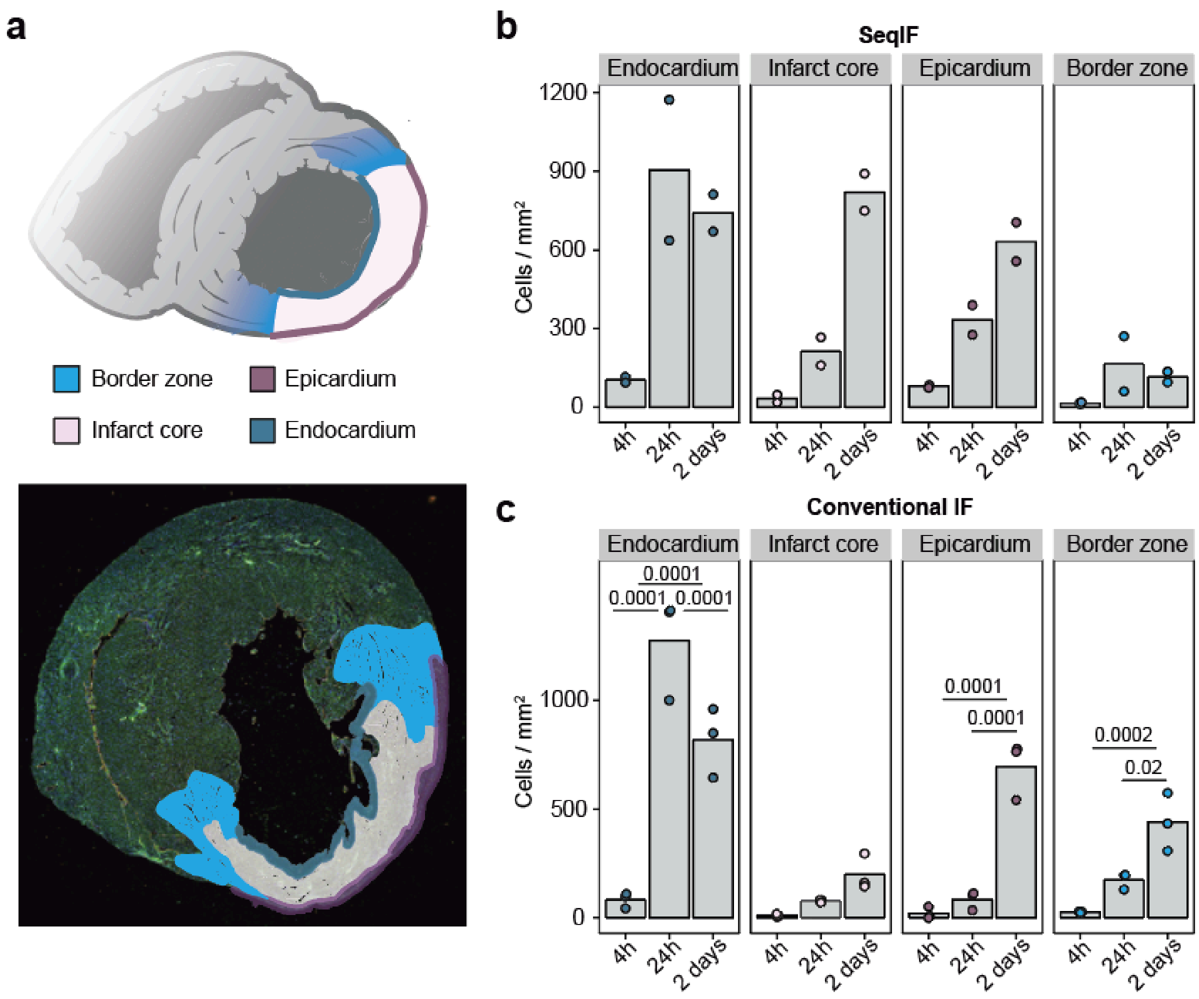


**Supplementary Figure 7: Quantification of immune cell infiltration from SeqIF and conventional IF imaging. a)** Schematic highlighting different regions for quantification of immune cell infiltration (top). Representative staining of WGA (green), CD31 (yellow) and DAPI (blue) with annotated regions 24h post-MI. **b)** Quantification of CCR2+ Mo/Mϕ for biological duplicate SeqIF images in different regions as depicted in a. **c)** Relative numbers of CCR2+/CD68+ Mo/Mϕ in different regions of the heart as depicted in a, using conventional immunofluorescence staining for CCR2, CD68, CD31, WGA and DAPI. Bars show mean abundance and points represent individual measurements. *P* values were determined by 2-way ANOVA followed by Tukey's multiple-comparison test. Only significant comparisons between timepoints within each region are displayed.

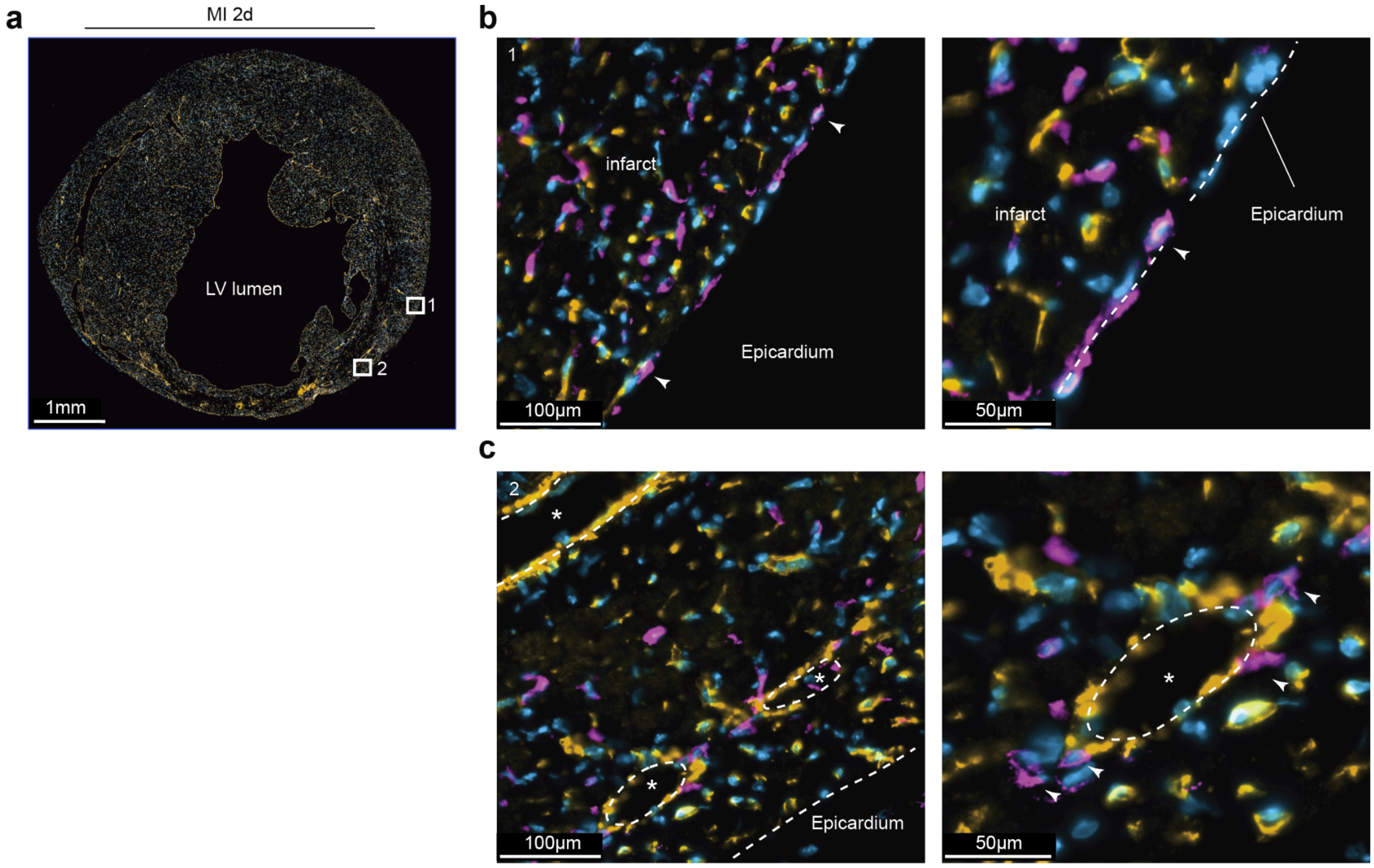


**Supplementary Figure 8: CCR2+ cells in proximity to epicardium and epicardial vessels. a)** Overview section 2d post-MI with staining of CD31 (yellow), CCR2 (magenta) and DAPI (blue). **b)** and **c)** Magnified areas with a focus on the epicardial region displaying CCR2+ cells on the surface of the epicardium (arrows) and epicardial vessels (marked by asterisks). The mid and right panel displays magnifications of the marked box in the left overview panel.

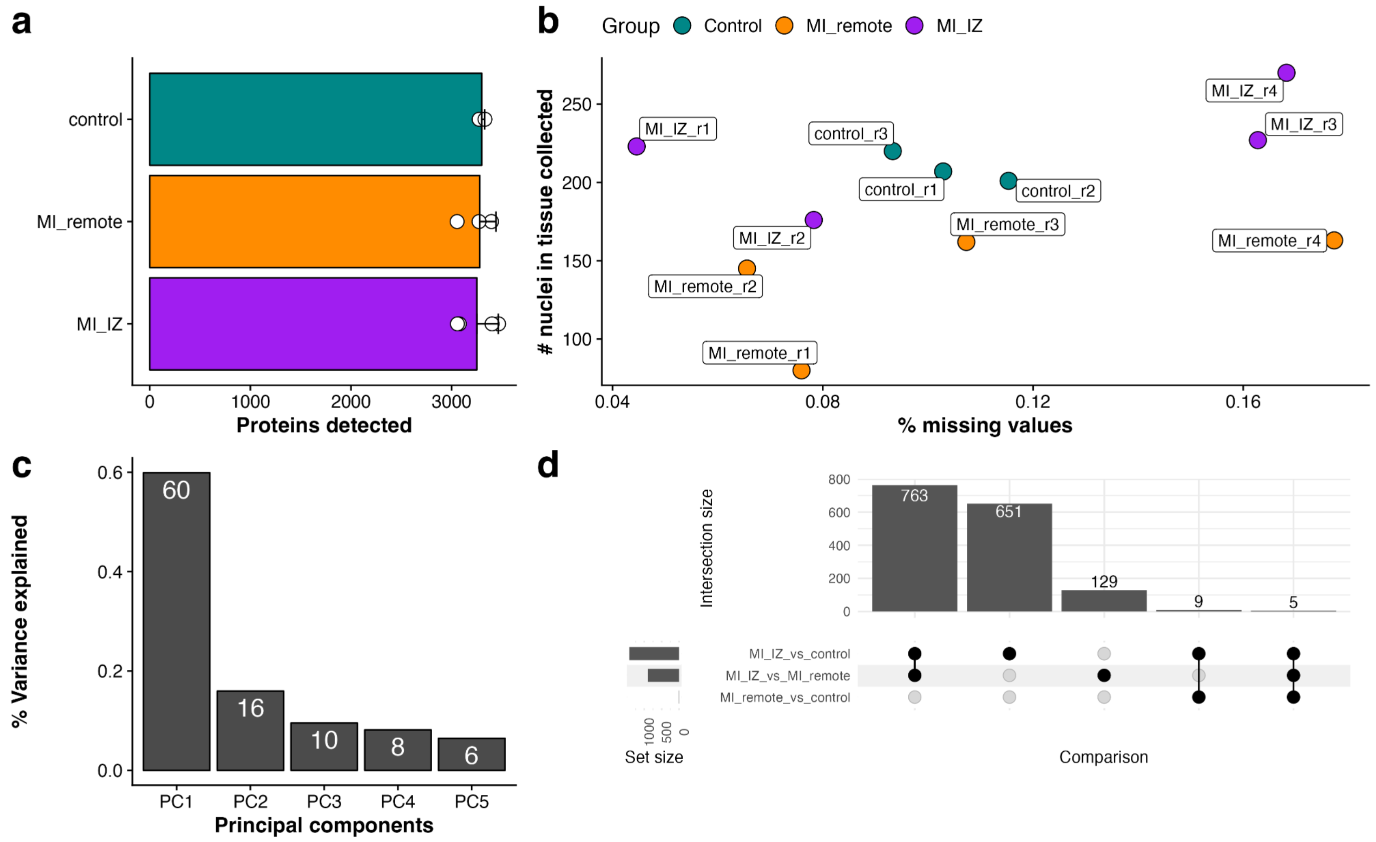


**Supplementary Figure 9: Quality control of DVP data. a)** Number of proteins identified per sample within each treatment / subregion group. **b)** Scatterplot showing the percentage of missing values in the proteomic data against the number of nuclei that were detected within the region marked for laser microdissection **c)** Percent variance explained by the first five principal components for proteomics data. Principal component one separates samples by treatment group / subregion and explains 60% of the variance within the data. **d)** Upset plot showing overlap between significantly differentially expressed proteins between all three laser microdissected regions.

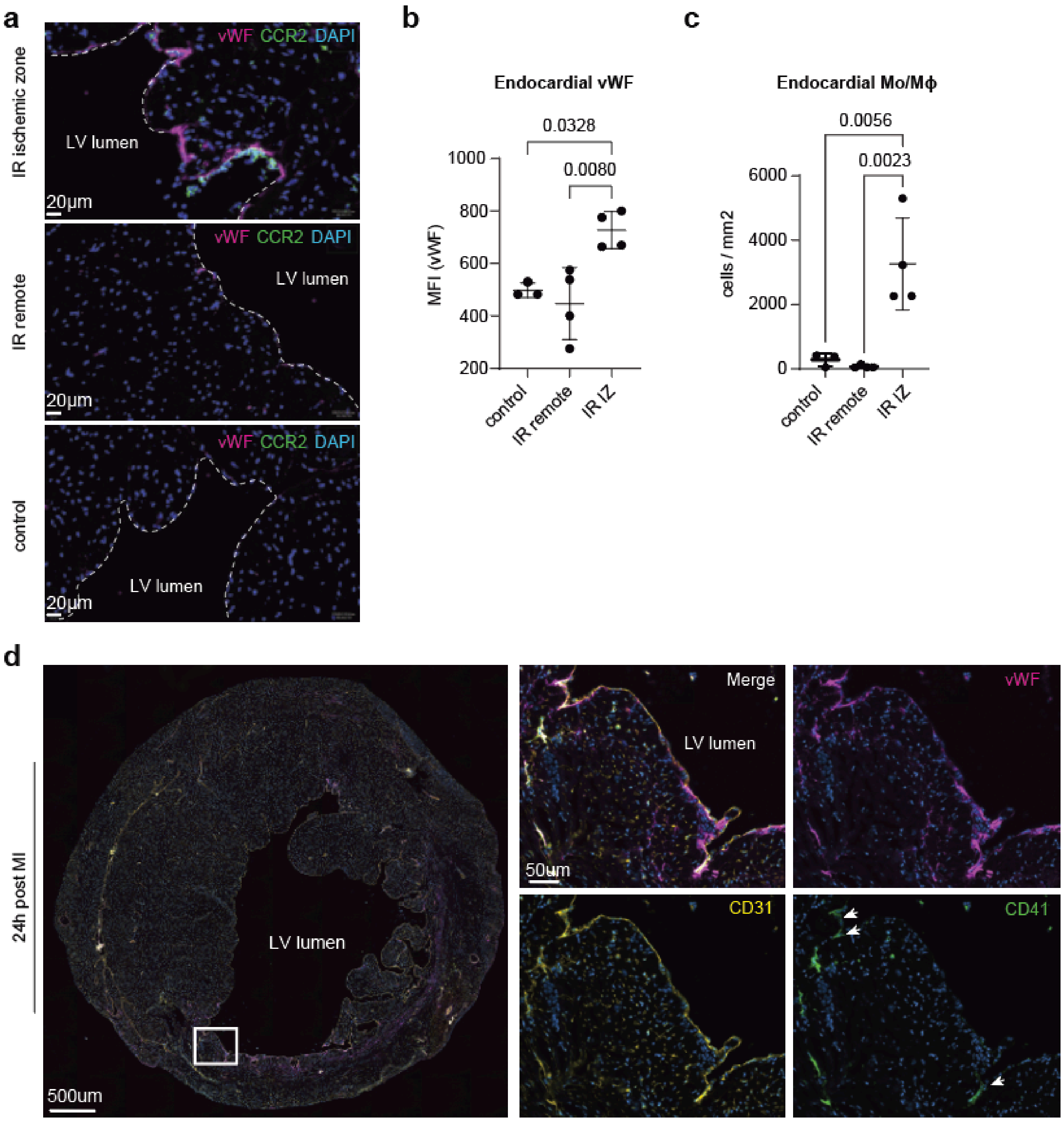


**Supplementary Figure 10: Immunofluorescent staining of CCR2+ Mo/Mϕ and endocardial vWF after ischemia/reperfusion injury. a)** Immunofluorescence stainings of CCR2 and vWF in the infarct zone 24h after ischemia/reperfusion injury. **b)** Quantification of endocardial vWF after ischemia/reperfusion injury. **c)** Quantification of infiltrating Mo/Mϕ at the endocardial region in both control hearts and hearts after ischemia/reperfusion injury. **d)** Co-staining of vWF, CD31 and CD41 in the infarct region 24h after permanent occlusion indicates a platelet-independent mechanism. Arrows indicate CD41+ platelets in some but not all vWF+ areas.
